# Mechanistic Insights into the Selective Targeting of MLX to Triacylglycerol-Rich Lipid Droplets

**DOI:** 10.1101/2024.07.15.603605

**Authors:** R. Jay Braun, Jessica M. J. Swanson

## Abstract

The activation of transcription factor Max-Like Protein x (MLX) is modulated by competition between active dimerization and inactive association with cytosolic lipid droplets (LDs). However, LD association has been shown to depend on the neutral lipid composition. This work explores the mechanism by which MLX specifically targets LDs rich in triacylglycerol (TG) over those with abundant sterol esters (SE). We compare the association ensembles for a potential minimal targeting sequence, an amphipathic helix-loop-helix hairpin, and the full dimerization and cytoplasmic localization domain (DCD), finding the latter requires larger packing defects and quantifiably alters LD membrane properties. Surprisingly, direct interactions with TG neutral lipids are not observed for either sequence. Instead, targeting to SE-rich LDs is blocked for both sequences by insufficient packing defects. We additionally explore the full mechanism of hairpin association, aiming to understand sequence-specific features that enable strong membrane association. We find that there are multiple association pathways, but that each involves a catch, dive, snorkeling, and embedding phase. The combination of multiple catch and dive residues placed on opposing ends of amphipathic helices lengthens the catch phase, greatly enhancing association in a manner that resembles kinetic selection. Once bound, locking interhelical interactions block dissociation. Collectively, our findings suggest that in addition to relative binding affinities, both kinetics and altered surface properties due to protein association could influence competition within the LD proteome.

**Significance:** The transcription factor Max-Like Protein x (MLX plays) a central role in metabolic regulation by responding to nutrient status and, simultaneously, neutral lipid composition. This work reveals how MLX selectively targets triacylglycerol-rich lipid droplets (LDs) through sequence-specific interactions with packing defects. We show that LD surface modulates MLX binding and that MLX in turn alters monolayer properties, highlighting a dynamic interplay between protein association and membrane properties. These findings provide new insight into how protein localization and function may be regulated at LD surfaces, with implications for nutrient sensing and, more broadly, transcriptional control relevant to metabolic and disease states.

## INTRODUCTION

Lipid Droplets (LDs) are cellular organelles that control lipid storage, distribution, and processing. In addition to their expected roles in lipolysis, lipogenesis, and the production of membrane building blocks, LDs have been found to play increasingly diverse roles (1–5), including in transcriptional regulation (6). Max like protein X (MLX) (Figure 1, S1a) was the first transcription factor identified to associate with LDs (6). As a member of the MYC- and MAX-centered network of transcription factors, MLX forms a dimer using a basic helix-loop-helix-leucine architecture (Figure 1a) that can bind DNA and influence expression levels. Unlike MYC and MAX, MLX has a small dimerization and cytoplasmic localization domain (DCD, Figure 1b) that regulates its activity. When bound to one of its dimerization partners (another MLX in a homodimer (Figure 1a-b) or MondoA or ChREBP as a heterodimer (Figure 1c-d)) (7–9) MLX can translocate to the nucleus where it regulates the transcription of proteins involved in glucose metabolism (10–15). In the presence of LDs, however, MLX can be stalled in the cytosol for sustained periods due to the DCD associating with LD surfaces with a slow apparent off-rate (6). In this manner LD association alters MLX-driven gene expression, providing a link between lipid storage organelles and glucose metabolic regulation. This work explores the mechanism by which MLX selectively associates with a subset of LDs.

**Figure 1.**
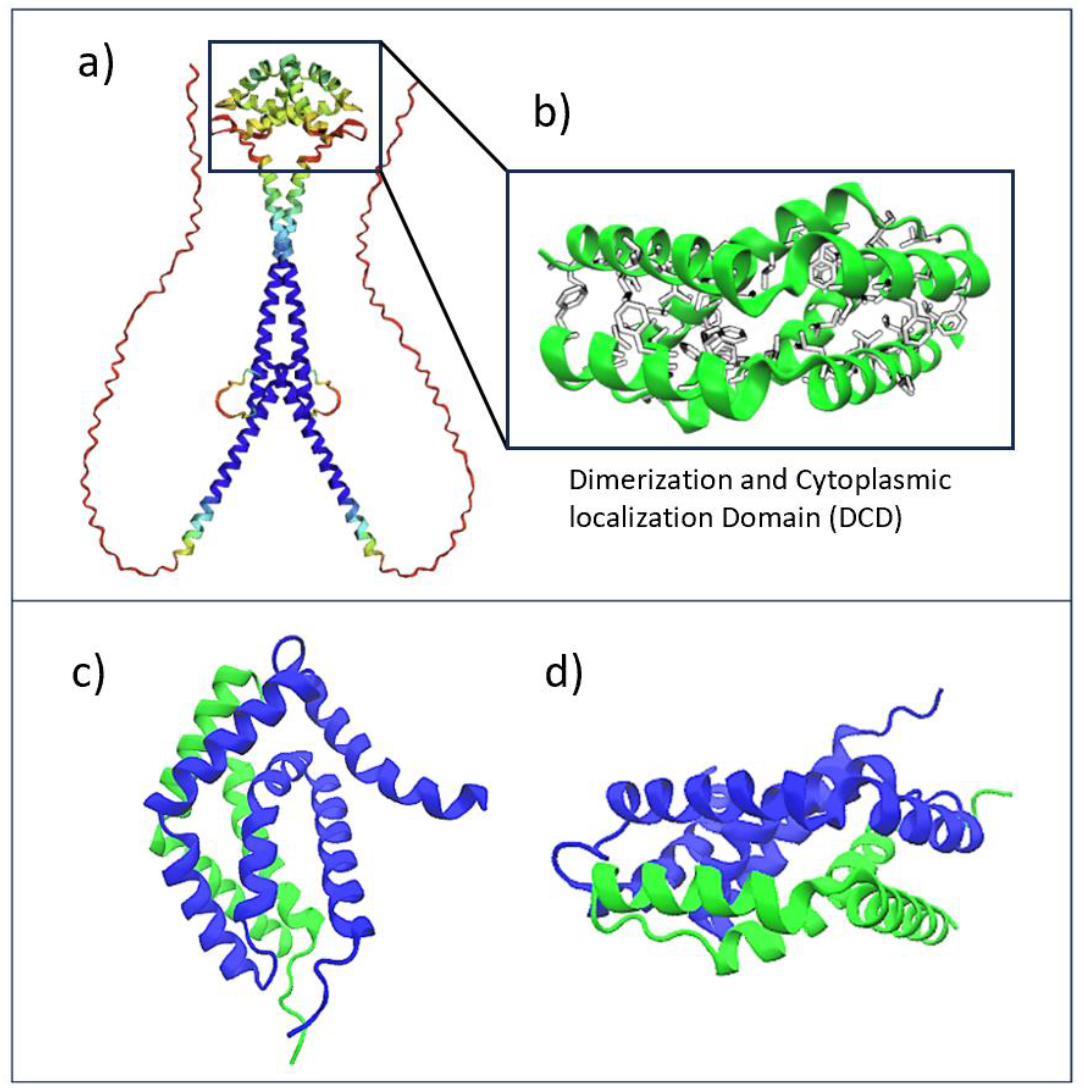
The predicted structures of a) the MLX homodimer, b) the homodimeric DCD, and the heterodimeric MLX-MondoA DCD from the c) top and d) side views.

LDs are uniquely characterized by a phospholipid (PL) monolayer membrane, unlike the bilayer membranes found surrounding other organelles. This monolayer encapsulates a neutral lipid core primarily composed of triacylglycerols (TGs) and sterol-esters (SEs). Decorating the monolayer is a dynamic array of proteins, collectively known as the LD proteome, which facilitate diverse lipid-related processes (1, 2, 4, 5, 16, 17). Understanding how proteins selectively associate with LDs at different stages of their lifecycle is an emerging area of LD research(2, 18, 19). Proteins that migrate from the cytosol, traditionally referred to as class II and now termed cytosol to LD (CYToLD) proteins (2), typically target the LD monolayer via amphipathic helices (AHs). These helices embed into the numerous large packing defects on the LD monolayer, where hydrophobic residues interact with the membrane’s exposed hydrophobic PL-acyl tail or neutral lipid regions. Meanwhile, polar residues engage with water or polar PL-headgroups. Another group of proteins, known as class I or endoplasmic reticulum (ER) to LD (ERtoLD) proteins, translocate from the ER membrane to the LD monolayer through membrane bridges. These proteins often feature membrane-embedded motifs, such as hairpins or longer U-shaped structures (2, 20, 21). In some cases, these motifs can possess polar or charged residues, which interact with ER-PL headgroups before engaging with the LD upon translocation. Together, these distinct protein classes regulate LD dynamics, including lipolysis, lipogenesis, and lipid trafficking—key facets of cellular lipid homeostasis.

LD proteomes have been shown to change in response to neutral lipid composition, with high TG or SE-compositions identified in distinct LD subpopulations (22–28). Although the physical origin of proteome remodeling is not fully understood, it has been proposed that the selective protein targeting to subpopulations is at least partially driven by the unique physical features of TG-vs SE-rich LDs (22). TG-rich LDs feature an amorphous core in which TG tails intercalate with the LD monolayer membrane, leading to larger and more frequent membrane packing defects (regions where the PL headgroups splay apart to expose hydrophobic acyl tails to water) (20, 24, 29). Additionally, TG molecules can intercalate into the PL monolayer and adopt conformations similar to PL molecules with their glycerol group facing the cytosol and their acyl tails packing in with PL tails (24). These surface-active TGs (SURF-TGs) explain the ∼15% increase in area per lipid headgroup measured experimentally (28, 30) in LD monolayers compared to the ER bilayer, their site of origin with the same PL composition. TG-rich LDs additionally create chemically distinct packing defects and, thus, LD-specific handles for proteins to target (2, 25, 31). SE-rich LDs differ substantially from their TG-rich counterparts, in part due to their propensity to undergo a phase transformation when SE:TG ratios are sufficiently high (26, 32). Under certain conditions (e.g., stress or starvation), high SE concentrations can transition into a liquid-crystalline smectic phase characterized by the formation of concentric rings beneath the PL monolayer (6, 23, 26, 32). Simulations have shown that this transformation increases the internal ordering of the droplet core and propagates outward to the monolayer, reducing the overall number of large packing defects by ∼50% (22). Furthermore, the denser PL packing expels SURF-TGs, thereby limiting opportunities for certain proteins to bind. Collectively, these changes lead to a reprogramming of the LD proteome, as some proteins are displaced while others are newly recruited, ultimately altering cellular and metabolic pathways (23, 33).

Herein we use a suite of computational approaches to investigate how MLX associates with lipid droplets, focusing first on a short double-helical region (residues 228–244) referred to as the MLX hairpin, which serves as a minimal model to uncover mechanisms of membrane binding and LD-core dependent specificity. Unbiased simulations show that the hairpin selectively associates with TG-rich LDs but not SE-rich LDs, capturing the preference for specific LD subpopulations. These simulations reveal that conserved hydrophobic and charged residues drive association through a catch, dive, anchor, and embed mechanism, and that interhelical contacts stabilize the bound conformation. We then examine the full DCD, which adopts a conformation nearly identical to its dimerized form when bound to TG-rich monolayers, but a different conformation in solution, suggesting a structural basis for toggling between membrane binding and dimerization. Interestingly, MLX targets shallow PL-acyl tail–induced defects, differing from the deeper insertion strategies used by other LD-targeting proteins (25, 34). Free energy of association calculations confirm that both the hairpin and DCD bind favorably to TG-rich droplets while being excluded from SE-rich systems. Finally, we show that MLX binding alters the physical properties of the LD monolayer, reducing packing defects, displacing SURF-TG molecules, increasing tail order at intermediate distances, and lowering lipid mobility. Together, these findings demonstrate how amphipathic helices achieve selective targeting and how even small proteins can reshape the surface properties of lipid droplets.

## METHODS

### Sequences and Structures

The association mechanisms of two MLX sequences were studied: the MLX-hairpin including residues 208-244 (Figure 2a) and the longer DCD including residues 177-244 (Figure 2b). These sequences were chosen because the MLX-hairpin was initially thought to be the shortest binding sequence. However, subsequent experiments revealed that it may also bind the ER bilayer (R. Farese and T. Walther, personal communications, 2022). In contrast, MLX-DCD has been shown to bind specifically to TG-rich LDs (6). Thus, these two segments provide a rich comparison to better understand selective targeting to ER-like membranes and TG-rich LDs (hairpin) versus specific targeting to only TG-rich LDs (DCD).

**Figure 2.**
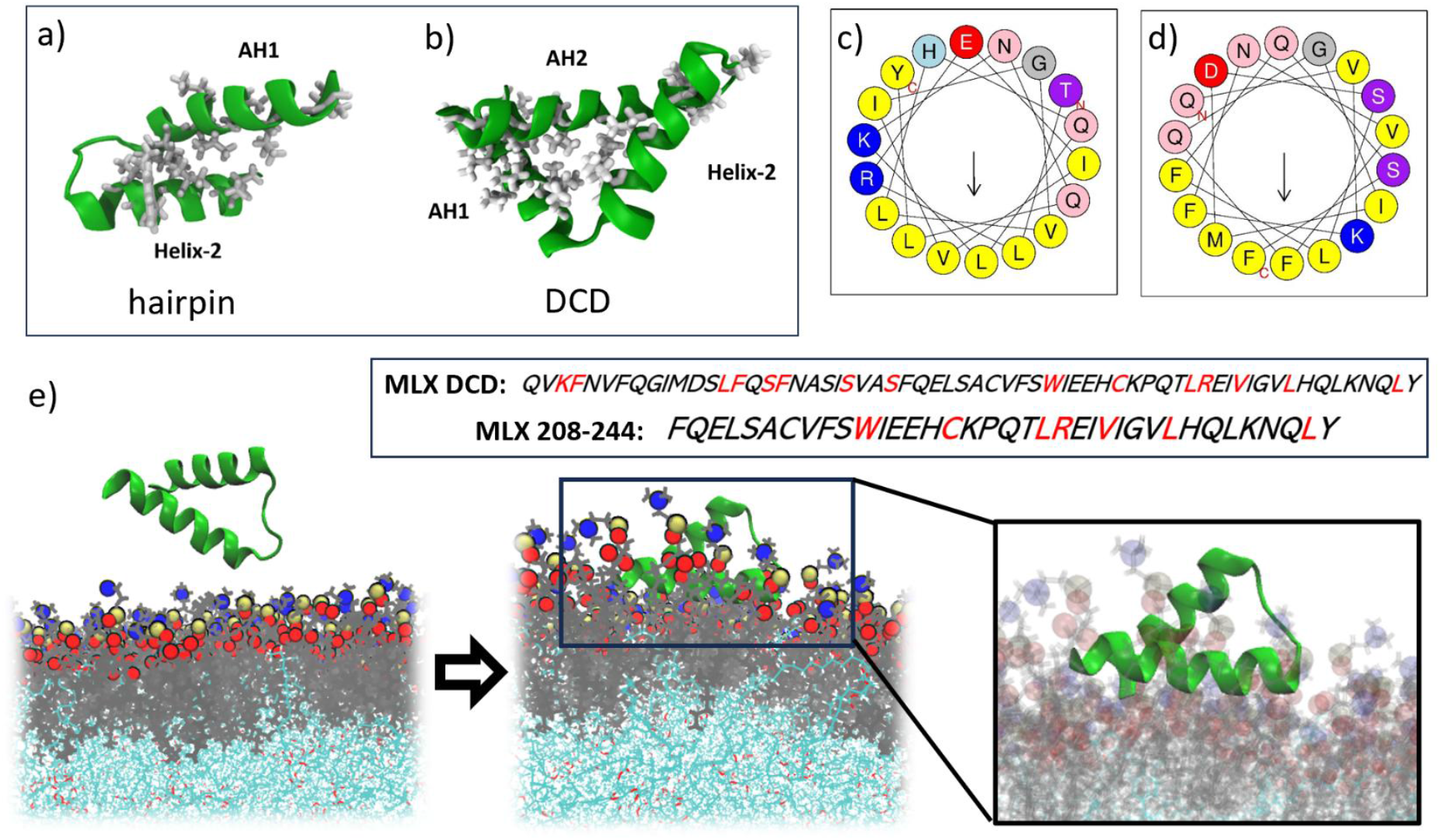
a) Predicted structures of MLX-hairpin (208-244) and (b) MLX-DCD (177-244). Conserved residues are highlighted in red in the written sequences below. c) Both sequences share a conserved AH1 that contains multiple Leu, Val, and Ile residues. d) MLX-DCD has a second AH2 with several Phe residues. e) Simulation structures of the initial cytosolic (left, water not shown for clarity) and LD-bound (right) MLX-hairpin.

In the absence of experimental structural information, multiple tools were used to predict both structures, including Rosetta Ab Initio, AlphaFold2, and RoseTTAFold. Virtually identical structures were predicted (see *Supplementary Methods* for full details). The MLX-hairpin contains two short helices (Figure 2a), one of which is clearly amphipathic (AH1) (Figure 2c) with multiple conserved residues (colored red in written sequences in Fig. 2: Leu228, Arg229, Val232, Leu236, and Leu2430) and several additional hydrophobic residues (Ile231, Val235, and Leu239). The larger MLX-DCD has an additional AH (AH2) (Figure 2d) that precedes helix-2 in sequence (Figure 2b). Although the DCD contains the hairpin and the two sequences have the same residues involved in AH1, helix 2, and the first kink, they take on different tertiary structures when solvated (Figure 2a-b). This is expected given the different hydrophobic residues that would naturally bury in an aqueous environment. The structures are further supported by the stability of the helices and kink locations throughout the simulations described below. Thus, the AHs are not unstructured in the cytosol, as is the case for some CYToLD proteins (2, 25, 29).

### Unbiased MD to capture real dynamics of association for the MLX-hairpin

#### Simulation Protocol

To explore the dynamics of membrane association, we first conducted long-timescale simulations of the shorter MLX-hairpin interacting with 3 LDs, each with a different neutral lipid composition: pure TG, 50:50 SE:TG, and 90:10 SE:TG (representing a SE-rich LD). The representative LDs were equilibrated in long-timescale simulations from our previous work (22, 24) using the CHARMM36m forcefield (35, 36). Our choice to use this forcefield is based on recent validation for LD monolayer membranes (37). Previous work has highlighted limitations in the CHARMM36 lipid forcefield (also referred to as CHARMM36-standard) for TG, as the glycerol backbone is overly hydrophilic in the LD core, leading to an underestimated surface tension at the water-TG interface and overhydration of the LD core relative to the hydration of bulk olive oil (38, 39). Alternative models, such as CHARMM36-cutoff (38) and CHARMM36/LJPME-r (40), address these issues by reducing the charges to a much more hydrophobic glycerol group until the experimental water-TG interfacial tension is matched. However, these changes also prevent SURF-TGs from migrating into the LD monolayer and hence fail to capture the experimentally measured monolayer expansion of ∼15% relative to a bilayer of the same composition (28, 30, 31). This discrepancy has raised concerns about both the occurrence of SURF-TGs and the overall validity of the CHARMM36-standard model. However, recent work using the Drude2023 lipid forcefield for TG demonstrated that enabling a dynamic charge distribution that responds to polar interface and hydrophobic LD core environments recapitulates all experimental observables, with naturally occurring SURF-TG levels and LD packing defects closely resembling those produced by CHARMM36-standard and quite different from CHARMM36-cutoff or CHARMM36/LJPME-r (37). Since MLX interacts with the LD monolayer, we opted to continue with the CHARMM36-standard model to ensure proper representation of the interfacial region, featuring SURF-TGs and Drude-consistent packing defects. It should be noted that for proteins deeply embedded within the LD core, it is not yet clear how TG polarization influences protein interactions.

The LDs were built, as previously described (22, 24, 25), using trilayer models, which remain essential for computational efficiency at the all-atom resolution. In these trilayers, a neutral lipid core is inserted between the two leaflets of a phospholipid bilayer. Put briefly, bilayers with an ER-like composition (88:37:10 POPC:DOPE:SAPI) (41) were built using CHARMM-GUI (42–44). Neutral lipid cores with varying SE:TG ratios (pure-TG, 50:50, 90:10) were pre-equilibrated for 500 ns in NPT at 310 K. Leaflets from the bilayer were separated and placed around the neutral lipid box to create monolayers, solvated with 50 Å TIP3P (45) water and 0.15 M NaCl on both sides. Final systems were simulated in GROMACS 2019.4 (46) for 8 µs with a 2 fs timestep using PME (47) for electrostatics (1.0 nm cutoff), LJ cutoffs from 8–12 Å (force-switch), and semi-isotropic Parrinello-Rahman (48) pressure coupling. Temperature was maintained at 310 K via Nose-Hoover (τ = 1 ps) (49, 50), and bonds to hydrogen were constrained using LINCS (53). Initial energy minimization (steepest descent, 5000 steps) removed TG–monolayer contacts. Trajectories were extracted every 100 ps. For a more detailed description of the simulation set-up and long equilibration, please visit our previous work (22, 24). Note that an 8 nm LD core was selected in this work. Although previous coarse-grained simulations have demonstrated some degree of long-range order of mixed SE and TG LD cores up to 8 nm (51), our previous work showed no difference in LD monolayer properties between an 8 nm and 16 nm cores (24). Since our aim was to capture sufficient sampling of a slow association process, the 8 nm core was selected for computational efficiency. A representation of the LD trilayer system is available in Figure S2.

In this work, the MLX sequences were initially placed at least 2 nm above the LD monolayer. An energy minimization of steepest-decent over 5000 steps was used to get rid of bad contacts between protein and water (noting the LD systems were already equilibrated from the previous studies). Production runs were conducted for a minimum of 1 µs. 15 replicas were run for all systems. Visualizations of the molecular systems were rendered with Visual Molecular Dynamics (VMD) (52).

### Association Ensemble Analyses

#### Contact analysis

To describe the binding process, we used MDAnalysis scripting to measure the center-of-mass (COM) depth of residue sidechains relative to the phospholipid headgroups’ phosphorus atoms. This was complemented by counting contacts (atoms within 3 Å) between phospholipid and TG defects and residue sidechains, enriching our understanding of molecular interactions. We calculated radial distribution functions to quantitatively assess dominant interactions using the MDAnalysis Python library (53).

#### Defect size below protein

To determine the size of the packing defects below the proteins, we calculated the convex hull area of the protein residue interactions with the defect pockets. Calculating the convex hull below the cutoff of the P atoms of the PLs allows us to determine the size, and depth of defects directly below the protein. The convex hull of a group of points S in n-dimensional space can be thought of as the smallest convex shape that encompasses all points within S. For *N* points *p*_*1*_,*…,p*_*N*_, the convex hull *C* is then given the expression:

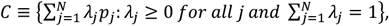

where *p* is a combination of 10 sidechains below the defect cutoff (54). If there was an inadequate number of sidechains below the cutoff, the convex hull was not counted (54). The size of the convex hull was averaged over the respective trajectory of 1000 frames. The MDAnalysis Python library was used for the scripting.

#### Packing defect analysis

We used a Cartesian-based algorithm to quantify lipid packing defects within monolayer leaflets, building upon established methodologies and algorithms (24, 29, 55, 56). Put briefly: For a monolayer leaflet, all lipid atoms whose positions were greater than the threshold (*Z*_*thr*_) were projected onto a 1 Å spacing two-dimensional (x–y) grid. The threshold (*Z*_*thr*_) was set to the average z-position of the phosphorous atoms of the leaflet minus 2 nm, which is sufficiently deep to cover the x-y plane. We first defined the van der Waals radii for a comprehensive set of atom types taken from the CHARMM36m (36) parameter set. This informed our overlap criteria between grid points and atoms, ensuring accurate defect localization. If a grid point overlaps with an atom (center of atom and grid point is less than atom’s radius plus half of the diagonal grid, 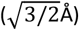, the z-position of the atom and the atom type are saved in the grid point. If the grid point overlaps with a polar PL headgroup, then it is not considered a defect. While scanning the grid using a depth-first search, if the grid point overlaps with PL-acyl groups or the LD-neutral lipids, then it is considered a defect. For each of these defect types, the neighboring elementary defects are clustered into one. If the clustered defect contains *N* elementary defects, it is considered to have a defect size of *N* Å^2^.

The probability of finding a defect with size *N* was computed and fit to an exponential decay function:

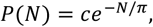

where *c* is the normalization constant and *π* is the packing defect constant. The packing defect constant represents how slowly the decay function falls off. A higher packing defect constant would generally mean that there is a higher probability of finding a larger defect. The original script for this packing defect analysis can be found at (https://github.com/ksy141/SMDAnalysis). We additionally modified the code to create cut-offs that excluded a desired radius from the protein once the protein associated to the respective leaflet of the LD monolayer. This allowed us to identify the alteration of monolayer defects due to protein interactions. To track the protein-defect interactions over time, we included further analysis to include the protein, while isolating specific heavy-atom interactions that were below defect threshold cut-offs. These protein atoms were included in the defect analysis and depth-first grid search but were not included as a defect type. If a heavy protein atom crossed the threshold of the defect in the x-y grid, it was counted as a defect-protein interaction. Modified packing defect script analysis scripts that account for protein cutoff distances, and identifications of top and bottom leaflets can be found at (https://github.com/swansongroup/PD). Error bars for packing defect constants were calculated using blocking averaging, with 10 blocks used over a 500 ns period.

#### Coordination analysis

The coordination of specific residues to the defects is determined by:

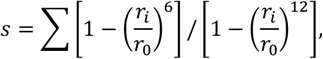

where *r*_*0*_ =0.4 nm, and *r*_*1*_ is the distance between the side chain group and defect group. Heavy atoms were used.

#### Tail Order Parameters

The tail order parameters are calculated with:

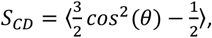

with *θ* being the angle between the position vector of a carbon atom of an acyl chain bonded to a hydrogen with respect to the system normal. A larger value of *S* Indicates increased ordering in the membrane. The brackets represent the ensemble average (57). The tail order parameters were calculated with MDAnalysis (53) scripting. We additionally calculated tail order parameters at specific radii cutoffs with respect to the protein. This code is available at https://github.com/swansongroup/surface_properties.

#### Mean squared displacement

To analyze the mobility of lipids in our system, we calculated the mean-squared-displacement (MSD) of lipid molecules over time. Lipids were tracked over the course of the simulation, and their positions at each time interval were recorded. The MSD was then computed using the Einstein formula through MDAnalysis (53, 58, 59).

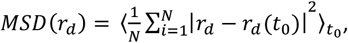

where *r*_*d*_ is the position of a lipid molecule at time *t*, and *r*_*d*_(*t*_0_) is its position at a reference time. Averaging was carried out over all lipids and over all choices of *t*_0_. By plotting MSD against time, we identified a linear regime corresponding to diffusive motion. The lipid diffusion coefficient *D* was then obtained from the slope of this regime using the two-dimensional relation *MSD* = 4*Dτ*, providing a quantitative measure of lipid mobility within the monolayer.

### Binding of MLX-DCD

Unbiased association of the DCD was first tested in four 1-µs simulations starting the solvent-equilibrated protein 2 nm above the LD monolayer. Since the three helices fold in upon each other to prevent the hydrophobic residues from interacting with the cytosol, no spontaneous association was observed on this timescale. This is not surprising given the expected kinetic barrier for the helices to open as they engage with the LD monolayer. To overcome this barrier and access the bound ensemble, a slight bias using steered MD (60) was used to open the AH1 and AH2 of the DCD to expose them to solution, which was subsequently removed. The COM of the protein moved towards the LD surface with no structural restraints in the z-coordinate until initial contact with the membrane surface was observed. A bias of 1000 kJ, with a rate of 0.001 nm/ps was used. Following this initial contact, regular production MD parameters were used as previously described. The simulation was run for an additional 3 µs, and the structural stability of the bound DCD was measured using root mean squared deviation (RMSD) (Figure S12). Additionally, we created a larger MLX DCD system (DCD-big) by doubling the *x and y* dimensions to 20 *nm* × 20 *nm* as well as a double-DCD system, with a DCD protein sequence on both top and bottom leaflets (Figure S3a-c). These systems were made to verify that the monolayer properties observed in the original simulations were not a consequence of size or pressure artifacts that might appear due to the DCD embedding into only one monolayer in a trilayer system.

### Quantifying association affinity

Umbrella sampling (US) (61) was used to determine the free energy of association for both the MLX-hairpin and MLX-DCD. The potential of mean force (PMF) was calculated as a function of the z-coordinate of the COM of the respective peptide sequence with respect to the distance from the average PL phosphorous z-coordinate plane. For MLX-hairpin, 22 windows were used, and for the MLX-DCD, 20 windows were used, with 0.125 nm used as the window spacing. The windows were obtained through natural binding in the pure-TG and 50:50 SE:TG systems, and through partial association event in the 90:10 SE:TG and for the DCD. Steered MD, as described in the previous section, was used to fully associate the proteins in which partial association occurred, to obtain initial structures for the bound windows. A harmonic potential with a force constant of 1000 kJ/mol nm^-2^ was used as a distance constraint. There was no helical restraint—the helices remained stable throughout the simulations. The rest of the simulation parameters were the same as described in the unbiased simulations above. Simulations were run for a total of 110 ns with the first 40ns discarded as equilibration ensuring conformational settling to each region of the reaction coordinate with a converged range of fluctuations. The Weighted Histogram Analysis Method (WHAM), specifically the tool g_wham (62), was used to calculate free energy profiles with uncertainties estimated using bootstrap resampling with 500 iterations (Figure S4). Autocorrelation analysis of the z-coordinate collective variable confirmed short integrated autocorrelation times (∼1 ns), indicating that the final 70 ns per window contained sufficient independent data (Figure S4c). Block averaging over 10 ns intervals and 7 blocks was also performed as a consistency check, yielding comparable uncertainty estimates (Figure S4a). Histogram overlap between adjacent windows was checked to ensure sufficient window spacings (Figure S4b). Simulations were run using GROMACS version 2019.4 (46).

## Results and Discussion

### MLX-hairpin targets TG-rich LDs and avoids SE-rich LDs through sequence-specific interactions

Similar to MLX, many CYtoLD proteins dynamically associate with LDs in response to metabolic shifts (2, 3, 5, 63, 64). Unlike other organelles like mitochondria and the ER, LDs lack protein-association machinery, so targeting is largely determined by protein-membrane interactions, relative association affinities, and relative protein/LD abundance (19, 65). The composition of neutral lipids in the LD core has also been shown to influence protein association (23). Our investigations begin with the MLX-hairpin, which was originally thought to be the shortest LD targeting sequence for MLX, but was later found to associate with ER-like membranes also (R. Farese and T. Walther, personal communications, 2022). Despite lacking a clear independent biological role, the hairpin serves as a useful model for understanding features that enable targeting to both ER and LD packing-rich environments while being excluded from SE-rich LDs. It also serves as a valuable comparison to the full MLX DCD, which only targets TG-rich LDs. Lastly, the smaller size and helical stability of the hairpin enabled us to explore for the first time the full association/dissociation mechanism of a CYToLD-like AH-containing protein.

To investigate the molecular basis of hairpin association, we performed parallel ≥1 µs unbiased simulations of the MLX-hairpin using the CHARMM36-standard force field, which was recently validated for LD monolayers as described in Methods (*Simulation Protocol*) (37). Each simulation began with the hairpin 2 nm above the surface (Figure 2e). Although observed properties are not expected to be statistically converged, unbiased simulations are essential for exploring the association mechanism with real dynamics unaltered by a Hamiltonian bias. Focusing on AH1 (residues 228–244), we defined full association as the center-of-mass COM of at least four hydrophobic residue sidechains descending to or below the PL phosphate plane (Figure 3a-b). This occurred in 8 of 15 simulations on pure TG LDs and 4 of 15 on 50:50 TG:SE LDs, but never on SE-rich (90:10) LDs (Figure 3c). Contact analysis demonstrating the interactions were primarily with PL acyl chains rather than TGs (Figure 3e-g, S5).

**Figure 3.**
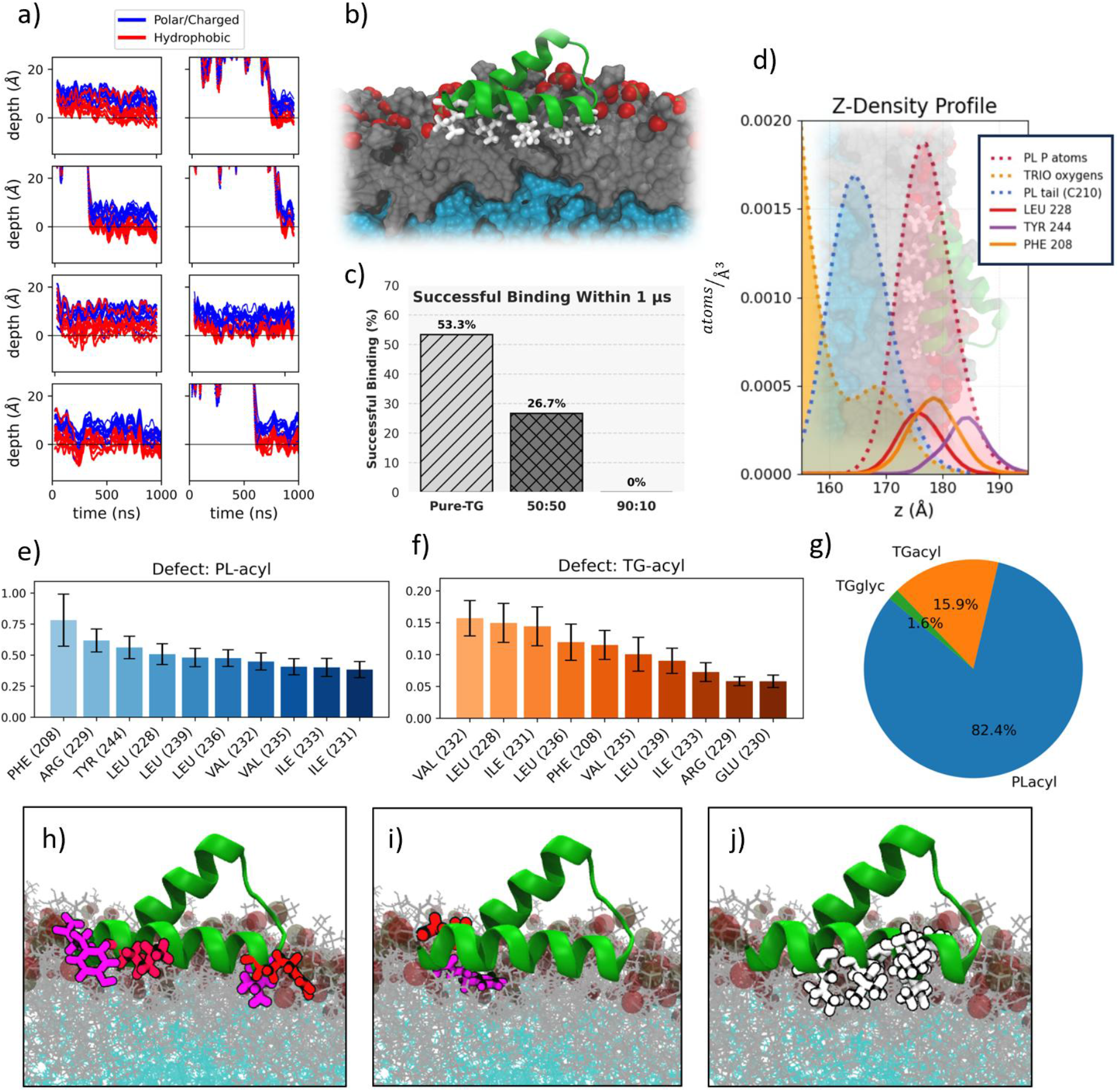
MLX-hairpin binds to TG-rich LDs via hydrophobic face. a) Time dependent depths of residues relative to the phosphate plane (black line) in 8 replicas differentiating the sequence of burial by hydrophobic (red) and polar (blue) residues. In all association cases, the hydrophobic residues are facing the membrane. b) Representative bound MLX-hairpin highlighting conserved hydrophobic residues (white) of the AH. c) The success rates for protein-LD binding. d) The depth of three residue’s COM are relatively shallow, right under the PL phosphorus (red) and interacting with the upper PL tail (blue). e) Normalized total defect interactions of both the PL-acyl and f) TG-acyl defects (error bars are standard-error-of-mean of all trajectories in which binding occurred). g) Interactions with PL-acyl defects are dominant. h) Catch residues Arg and Lys (red) interact with PL headgroups on opposite ends of AH1, while neighboring ‘dive’ residues Leu and Tyr (purple) dive into initial defect formation. i) A similar Glu/Phe catch-dive pair helps anchor the hairpin at the end of Helix-2. j) A series of leucines, valines, and isoleucines bury into defects to secure hairpin association.

To clarify how packing defects contribute to MLX association, we quantified residue interactions within a defined cutoff from membrane defects (see Methods, Movies 1 and 2). Surprisingly, the first three residues that interact with PL defects include Arg229 and Tyr244 (Figure 3e), a charged and polar residue expected to interact with PL headgroups or bury just below headgroups in the interfacial region, respectively—not to bury into a defect. This is explained by neighboring residues that create PL “catch” and defect “dive” duos on either side of AH1: Arg229/Leu228 and Lys240/Tyr244 (Figure 3h). Another such duo is found on the terminus of Helix-2: Glu210/Phe208 (Figure 3i).Notably, Arg229/Leu228 is conserved (Figure 2a) suggesting this catch-dive pair could play an essential LD association role in the full protein.

Between the anchoring catch-dive residues, a series of partially conserved hydrophobic residues, including leucines, valines and isoleucines, robustly interact with PL-acyl chains (Figure 3j), showing less interaction with TG-acyl defects or TG-glycerols (Figure 3e). Z density profiles (tracking the COM of three residues) confirm shallow membrane insertion near the PL headgroup region (Figure 3d). Although Lys240 does not show up in the dominant defect interactions, this is likely a consequence of the selected cutoff distance for defining packing defect interactions. Like Arg229, Lys240 works with Tyr244 in catching the opposing end of AH and itself contributes to stability by electrostatic interactions with PL headgroups (Figure S6a–c). Collectively, these findings demonstrate how specific residues recognize PL-acyl packing defects to selectively associate with ER-like membranes or TG-rich LDs, which contain robust defects.

It is worth noting that MLX’s association mechanism is distinct from other CYToLD proteins, like CCTα and CGI-58, despite all three featuring CYToLD AHs. CCTα binds to expanding LDs via Trp residues that interact with TG-glycerol defects. CGI-58, on the other hand, employs three Trp residues to penetrate deep within LD packing defects and into the LD core (25, 34). MLX sets itself apart, demonstrating a more versatile membrane-binding capacity by preferentially engaging shallow PL defects (Figure 3d-j), which could play a role in exchange between the LD surface and dimeric DCD interface, as further discussed below.

### MLX-hairpin association follows a heterogeneous catch-dive-anchor-embed mechanism with stabilizing conformational changes

Time-resolved defect-interaction analysis (Figure S7) reveals that the MLX-hairpin associates with the LD monolayer through multiple pathways, all culminating in a consistent bound pose. Consistent with the dominant interaction analysis above, key catch residues initiate membrane contact. Arg229, located at the hairpin kink, initiates binding in 5 of 6 tracked events (Figure S7a–f), likely due to its strong electrostatic interaction with PL headgroups and proximity to its partner dive residue, Leu228, which readily buries into initial defects (Figure 4a, S7a–b, e–f). Extending this role to the full protein could explain why this catch-dive duo is conserved (Figure 2a). In 3 of these cases, the catch-dive pair at the opposite end of AH1 (Lys240/Tyr244) briefly forms hydrogen bonds with PL headgroups before descending into PL-acyl defects (Figure 4a, S7a–b, f). Together, Arg229/Leu228 and Lys240/Tyr244 can anchor both ends of AH1 (Figure 4a). Other pathways include submersion initiated at the kink via SURF-TG (Figure S7c) or rapid hydrophobic insertion without a clear anchoring phase (Figure S7e). In one instance, Glu210/Phe208 located on the opposite face of helix 1 acts as the initial catch-dive duo (Figure 4a, S7d), highlighting variability in the starting point of association. Despite this variability, catch-dive residues are strategically positioned at opposing ends of the helices, enhancing the likelihood of successful engagement. Anchoring the other side of AH1 at the membrane interface further allows hydrophobic residues to transiently snorkel before full embedding (Figure 7a).

**Figure 4.**
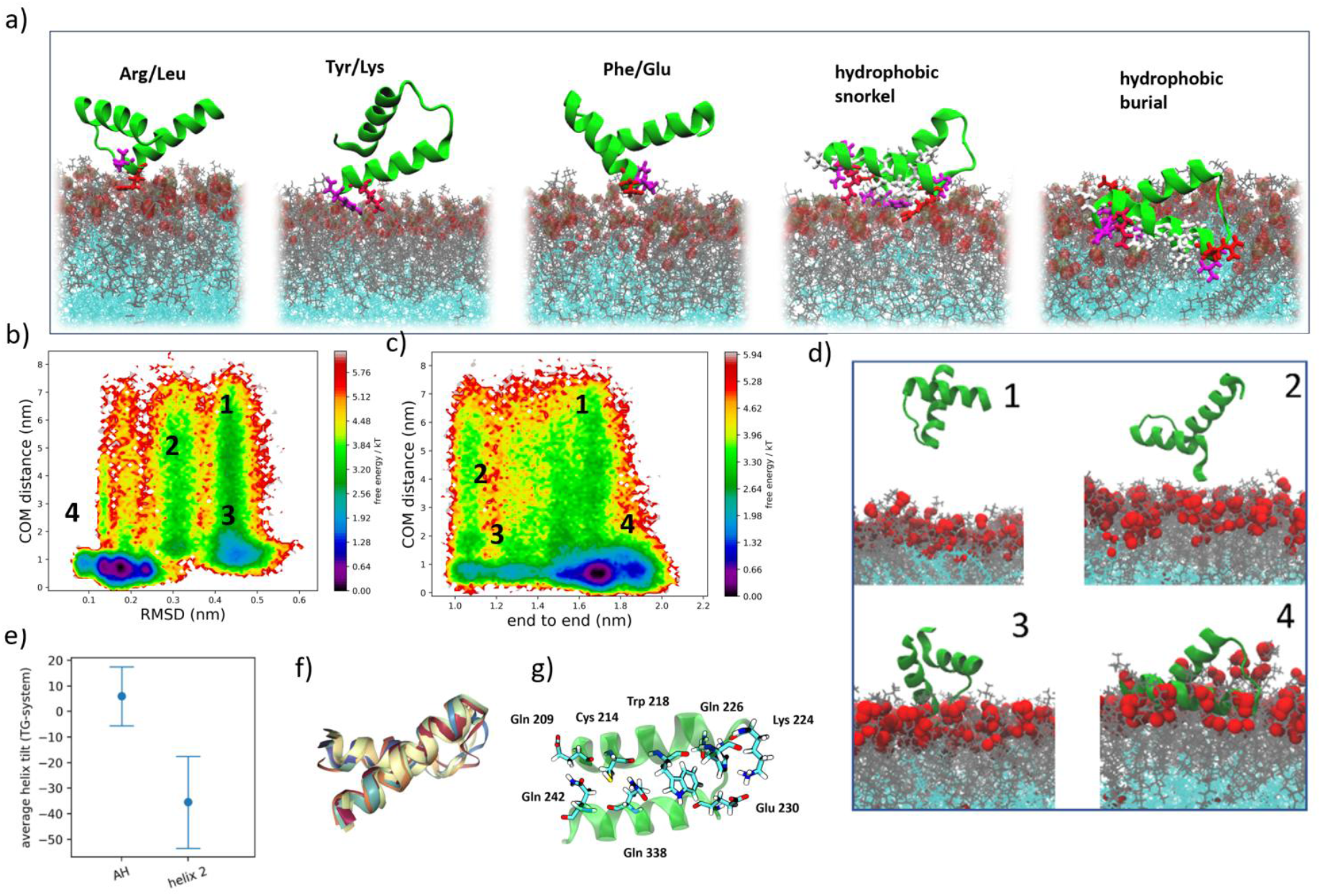
a) Several catch-dive pairs (Arg229/Leu228, Lys240/Tyr244, and Glu210/Phe208) initiate membrane association. After the initial catch and dive, hydrophobic residues snorkel on the surface, trying to embed into a defect. Finally, the hydrophobic residues find and create an adequate defect, burying the AH deep into the PL-acyl tail region. b) RMSD versus center-of-mass (COM) distance reveals multiple metastable wells, with the fourth well representing the most stable bound state. c) A similar trend is observed in the COM versus end-to-end distance plot, further supporting the identification of the stable conformation. d) Representative structures from each metastable well highlight the conformational diversity of the MLX-hairpin, with the fourth structure exhibiting the most compact and stable binding mode. e) The final bound pose maintains a specific tilt angle between the two helices and more narrow range relative to the membrane normal. f) Overlap of the bound conformation from the 8 trajectories that resulted in binding shows consistency in the final bound conformation, stabilized by g) multiple interhelical interactions.

The prevalence of this catch-dive phase suggests that kinetic factors influence selectivity, as proteins with longer-lived catch-dive states are more likely to fully associate once sufficient defects form. Similarly, monolayers with more frequent defect formation offer greater opportunity for fully embedding the caught protein. Multiple binding pathways enabled by distributed catch-dive residues increase the probability of stable anchoring, even if an initial contact is lost. Collectively, MLX exhibits a catch-dive-anchor-embed mechanism driven by flexible engagement routes and strategically placed residues (Movies 1, 2, 4).

In addition to interactions with the LD monolayer, the MLX-hairpin forms stabilizing interhelical contacts that reinforce the bound state. While AH1 and helix 2 remain independently folded throughout simulations, membrane association induces a clear conformational rearrangement. In the cytosol, the helices are flexible with variable tilt angles relative to the membrane normal (Figure S8a), but upon binding, they adopt a conserved crossed configuration: AH1 tilts upward by ∼5° and helix 2 downward by ∼35° relative to the membrane normal (Figure 4e–f, S8a). This geometry is stabilized by a network of hydrogen bonds, charged, and hydrophobic interactions, including new contacts between Lys224 and Glu230 near the kink (Figure 4g, S9c-d). The number of hydrogen bonds between the helices nearly doubles in the bound state (Figure 9b), and multiple leucine residues from each helix form a hydrophobic interface (Figure S9a, d). To quantify structural convergence, we projected the free energy landscape onto RMSD versus COM distance to the membrane and end-to-end distance versus COM distance to the membrane (Figure 4b–d). The bound ensemble consistently appears at low RMSD and a characteristic end-to-end distance when the protein is near the membrane. RMSD versus end-to-end distance alone is sufficient to delineate the bound state (Figure S8b), and alignment across eight independent replicas confirms convergence onto this pose (Figure 4f).

These interhelical interactions significantly contribute to the persistence of the bound state. As demonstrated for other systems (66), cooperative coupling between helices enhances membrane affinity by reducing the likelihood of dissociation, disengaging one helix requires overcoming stabilizing interactions with the other. This mechanism mirrors simulations observations in the SARS-CoV-2 fusion peptide, where interhelical interactions reinforced membrane binding (67). In MLX-hairpin, the defined tilt angles, converged structural ensemble, and strengthened interhelical network all point to a thermodynamically stable and kinetically persistent bound conformation. This cooperative effect, as explored for DCD below, could help explain MLX’s increased membrane affinity and slow off-rate (6), supporting the idea that interhelical stabilization is a key contributor to long-lived LD association. Although not biologically functional on its own, the MLX-hairpin illustrates a general strategy by which CYToLD proteins can achieve membrane targeting through catch-dive-anchor insertion and membrane-induced conformational locking.

### MLX-DCD binds large, shallow defects that mimic the dimer interface

Turning to the full DCD, it is helpful to have the context that the MAX and MYC protein families share a conserved basic leucine zipper architecture (Figure 1, S1), and recent NMR studies indicate that even the basic region retains partial helicity (67). Consistent with this, AlphaFold predicts high-confidence structures (pLDDT > 90) for the leucine zipper structure, as well as the DCD. Both the monomer and homodimer DCD structures adopt a triangular planar arrangement, in which AH1 and AH2 partially overlap and form stabilizing hydrophobic contacts (Figure 1, S1b). The predicted MLX-MondoA heterodimer shows a similar interface, with aligned DCD domains maximizing helical interactions (Figure 1b). These consistencies and the high pLDDT score give us increased confidence that the DCD structure used in our simulations is consistent with the MLX homodimer.

However, simulations of the isolated DCD monomer in solution show that without the dimer scaffold, the helices fold inward to shield hydrophobic residues (Figure 2b), forming a stable conformation that did not bind the LD in four 1 µs replicates. To explore the bound ensemble, a mild bias was therefore applied to open the DCD and expose the AHs, enabling stable membrane association (see Methods). Once bound, the DCD unfolded and inserted all three helices into the membrane, with AH1 and AH2 engaging PL-acyl defects (Figure 5a). The same key catch-dive, anchor, and embedding residues found in the hairpin stabilize the bound DCD, albeit with a significantly different interhelical arrangement (Figure 5a, S10). Additionally, four phenylalanine residues complement valines, leucine and isoleucine in AH2 engaging PL defects. Beyond these lipid interactions, the helices pack against each other in the membrane-bound state, forming stabilizing interhelical contacts that reinforce the inserted conformation. Such cooperative interactions may reduce the likelihood of spontaneous dissociation, consistent with the experimentally observed slow off-rate of DCD.

**Figure 5.**
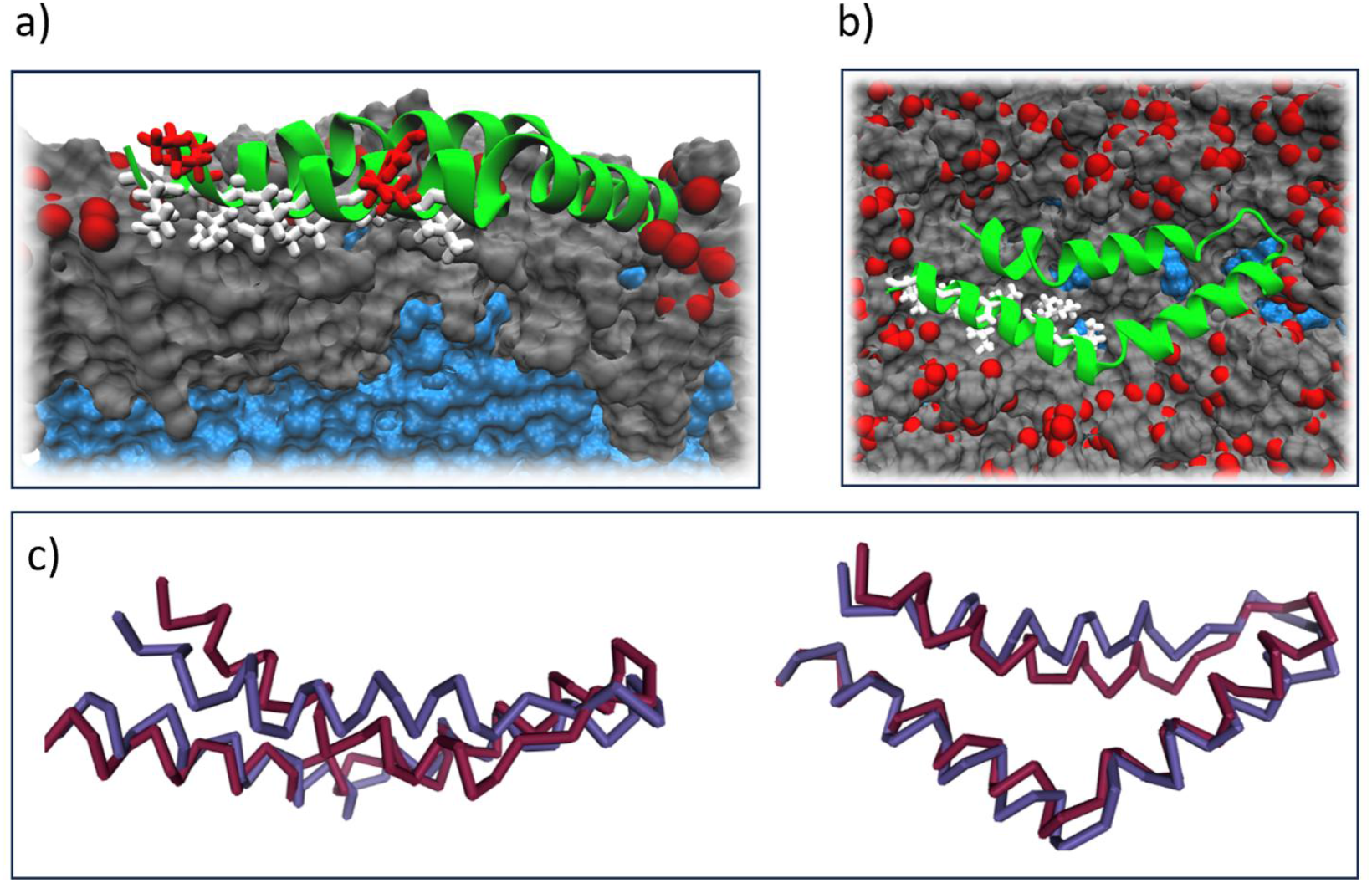
a) The DCD uses the same AH residues (colored in white) to bury into large, shallow PL-acyl defects. Arg229 and Lys240 still interact with PL headgroups. b) Top-down view of the DCD embedded into the PL-acyl defects. The structure is almost identical to its dimerized form. c) Nearly perfect overlap of one of the DCD homo-dimer chains (red) to the final LD-bound structure (blue).

Unexpectedly, the LD-bound DCD structure returns to closely mirrors its original dimer form, suggesting that hydrophobic burial at the dimer interface mimics that within the LD membrane (Figure 5b–c). This conformation remains highly stable once bound, as confirmed by backbone RMSD (Figure S12), suggesting that the DCD can toggle between dimerization and membrane association depending on cellular context. The compact mix of smaller hydrophobics (Leu, Val, Ile) on AH1 and bulky aromatics (Phe) on AH2 supports both dimer formation and stable LD binding. This shared interface may allow MLX to dynamically balance transcriptional activity and LD localization.

In contrast, other CYToLD proteins such as CCTα and CGI-58 rely on bulky Trp residues to penetrate deeply into LD defects (25, 34), pointing to a different recognition mode tuned to the deeper TG-induced membrane defects. Notably, DCD binding generates significantly larger packing defects than the hairpin: the average defect size across 1000 frames is 92 ± 17 Å^2^ for MLX-DCD versus 63 ± 16 Å^2^ for the hairpin (Figure S13). The larger footprint of the DCD both requires and induces more extensive defect formation, likely contributing to its selective affinity for TG-rich LDs, which have more frequent and larger defects than bilayers (22, 24).

These findings help explain the distinct modes of membrane association for the two MLX-derived sequences. The hairpin binds to membranes containing packing defects, including ER-like bilayers and TG-rich LD monolayers, but is excluded from SE-rich LD monolayers. It serves as a general model for how amphipathic helices engage defect-rich interfaces. In contrast, the DCD exhibits strict selectivity for TG-rich LDs, requiring larger or more persistent defects for stable insertion. Together, these results verify that MLX distinguishes between TG- and SE-rich LDs based on the physical properties of the LD monolayer, requiring sufficient packing defects for hydrophobic embedding, and demonstrate how modular amphipathic helices can support selective and distinct membrane targeting within the CYToLD family.

### Thermodynamics of association supports preferential targeting to TG-rich LDs

To evaluate whether insights from unbiased simulations reflect ensemble behavior and verify our simulations are consistent with experiment, we computed association free energy profiles for MLX-hairpin and MLX-DCD across LDs with varying compositions (pure TG, 50:50, and 90:10 CHYO:TG; Figure 6a–b). The MLX-hairpin exhibited strong affinity for the 50:50 and pure-TG LDs, with association free energies of approximately 10 to 11 kcal/mol. In contrast, the 90:10 system showed an unfavorable profile, with a 4 kcal/mol energetic preference for the cytosol. The MLX-DCD showed even stronger binding to TG-rich LDs, with a free energy drop of ∼19 kcal/mol, but was strongly repelled from the 90:10 monolayer, raising the free energy by ∼15 kcal/mol. These profiles support the conclusion that packing defect accessibility and monolayer density are key determinants of MLX-LD targeting. In the membrane-embedded state (z = 0 in Figure 6a), the MLX-hairpin consistently forms interactions between charged residues (Arg229, Lys240) and PL headgroups in both TG-rich and SE-rich systems (Figure S14). In the SE-rich LD, the PL tail defects remain inaccessible, preventing effective hydrophobic insertion (Figure S14, Movie 3), while for the TG-rich LD, the conserved hydrophobic residues can make plentiful interactions with these defects. The strong binding affinity of MLX-DCD likely reflects its larger size and greater membrane engagement. At 63 residues, the DCD induces packing defects averaging 92 Å^2^, a magnitude rarely observed in SE-rich LDs or bilayers (22, 24). Even the smaller MLX-hairpin incurs an energetic penalty in the 90:10 system due to the lack of available defects (Figure 6a, Movie 3). Together, these findings emphasize that the selective targeting of MLX is governed by the biophysical properties of TG-rich LD monolayers, particularly their ability to form large, shallow, PL-dominant packing defects

**Figure 6.**
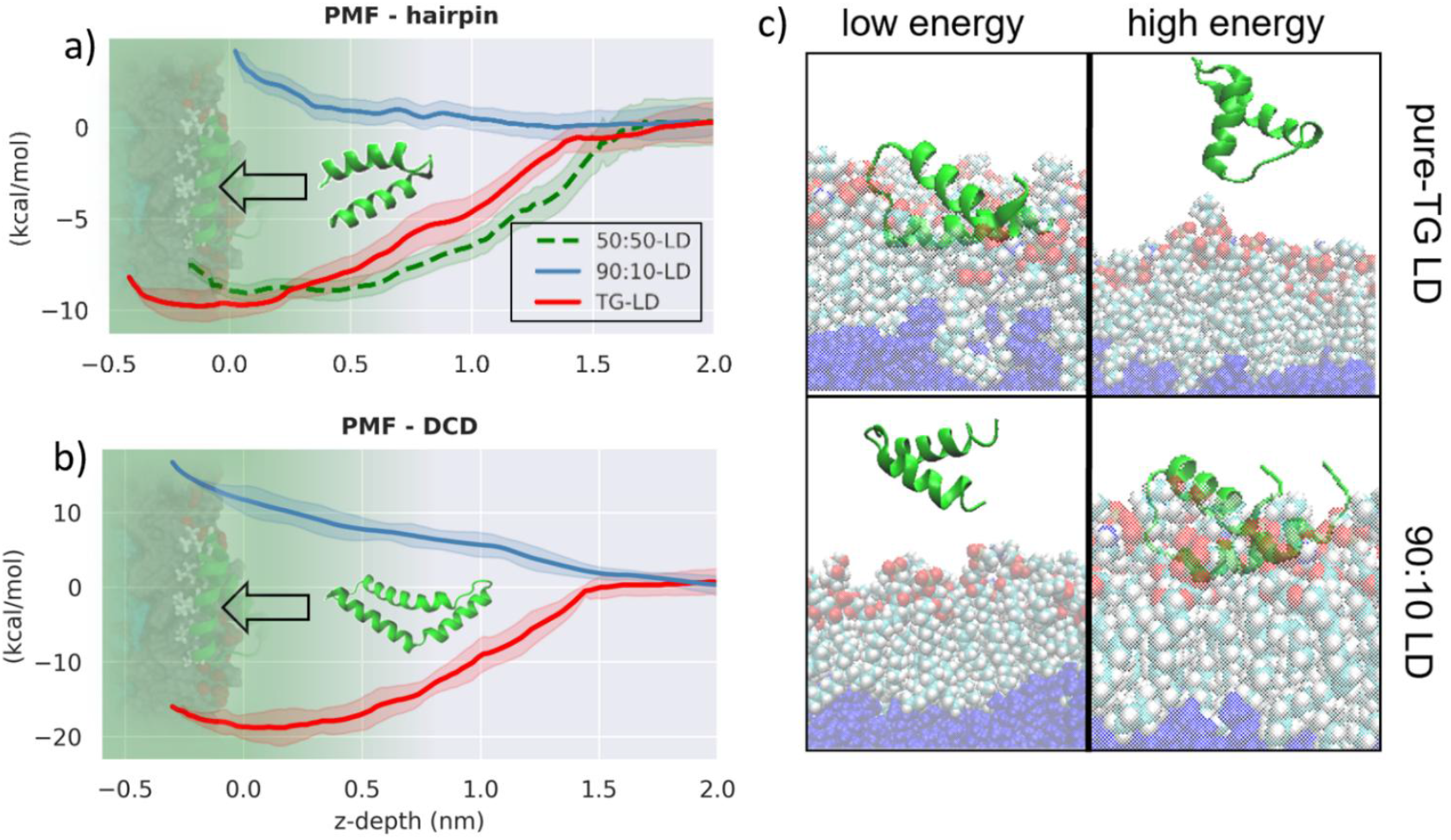
Free energy profiles for LD association of a) MLX-hairpin and b) MLX-DCD. c) The representation of low versus high energy conformations for the MLX-hairpin, emphasizing that the bound state is more stable for the pure-TG LD while the cytosolic state is more stable the SE-rich 90:10 LD. The sampled reaction coordinate (x-axis) for plots (a) and (b) is the z-COM distance of MLX from the phosphate plane (x = 0) of the LD monolayers. The green shaded area from -0.5 to 0.0 represents the PL-monolayer, while x > 0.0 moves into water. Note: The hairpin and DCD illustrations are not to scale with respect to the x-axis and are included solely for visual clarity.

### Association of MLX alters LD monolayer

An unexpected finding in this work was that the association of either the MLX-hairpin or MLX-DCD, as small as they are, appeared to alter the physical properties of the LD monolayer. To quantify this impact, we focused on monolayer defect organization and composition, PL-tail order parameters, and dynamics. To verify these properties are not artifacts due to the size of the system, we created the DCD-big system with twice as large *x* − *y* dimensions, quadrupling the surface area of the LD monolayer, and a DCD-double system with a protein inserted on both leaflets (see Methods).

#### Decreased Packing Defects

Upon MLX association, we observed a decrease in the probability and size of defects surrounding the protein on the protein-bound leaflet in each system (MLX-hairpin, MLX-DCD, DCD-big, DCD-double). This was quantified by measuring probability and likelihood of defects, specifically focusing on defects more than 1 nm from the protein. Defects are smaller, on average, on the protein bound leaflet, as shown visually in a top-down view of MLX-DCD (Figure 7a-b) and supported by the analysis of defect probability versus size (Figure 7d). We subsequently quantified the defect magnitudes by calculating the defect constants. A higher packing defect constant would generally mean that there is a higher probability of finding larger and more numerous defects (see Methods for details). All defect types were smaller on the protein-bound leaflets (Figure 7d, S15a-c). As expected, the magnitude of deviations decreases from the MLX-DCD to the MLX-DCD-big system (Figure S15c), reflecting that the protein’s influence is both spatially localized and diminished by the presence of additional free membrane area. This effect is more pronounced for PL defects, consistent with increased packing squeezing out SURF-TG molecules. However, the magnitude of deviations remains the same for the MLX-double system, verifying the pressure artifacts from the presence of a protein on only one leaflet are not the origin of the observed changes.

**Figure 7.**
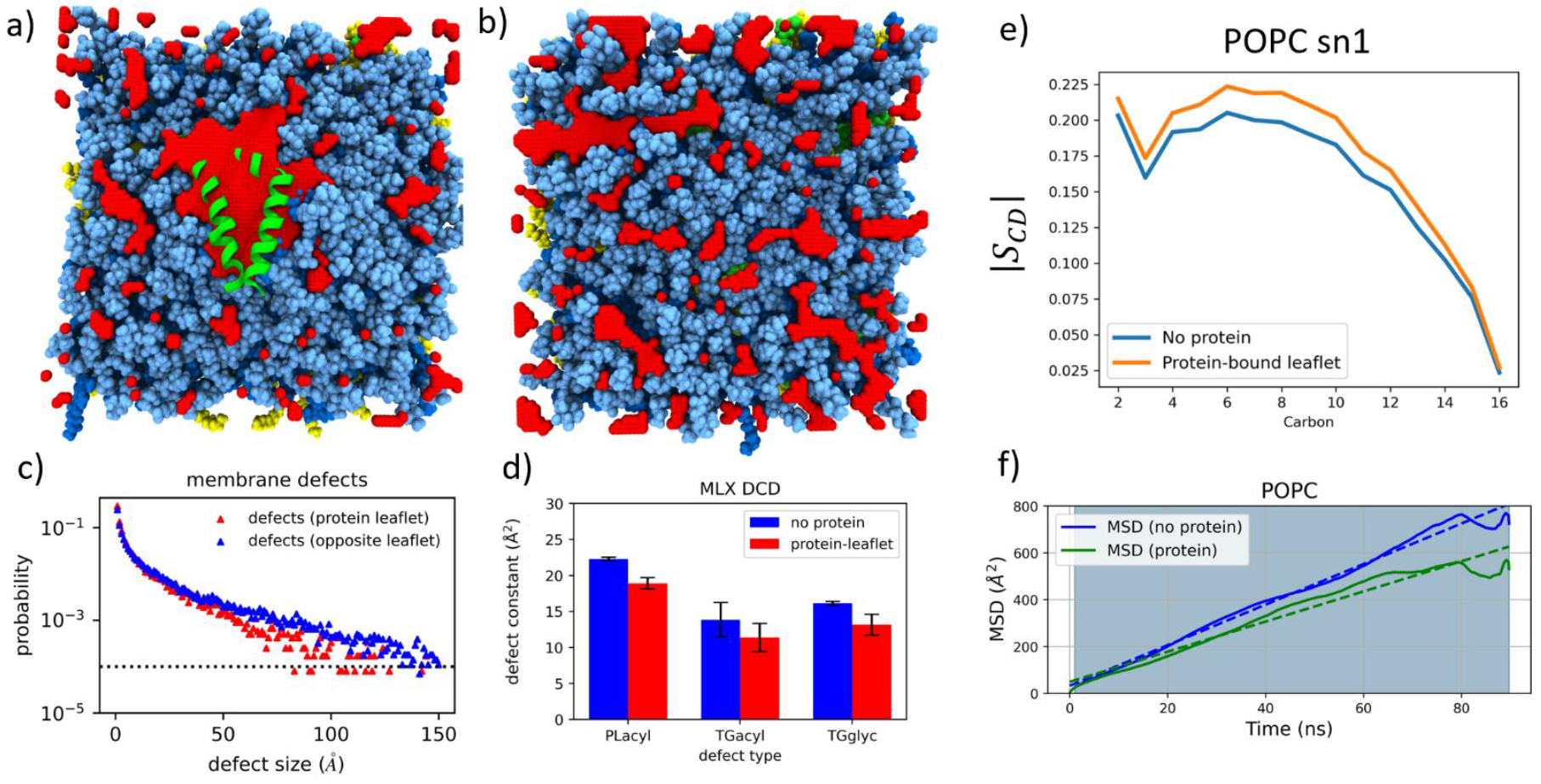
a) Packing defects in the DCD-bound leaflet, and b) LD monolayer without proteins, with red indicating defects and blue representing PLs. c) Defects size vs. probability for MLX-DCD demonstrates the protein-bound leaflet has smaller defects >1 nm from the protein. d) Packing defect constants, which measure the decay rate of the defect size vs. probability, demonstrate that all types of defects are smaller on the protein-bound leaflet. e) POPC sn1 tails are more ordered on protein-bound leaflets, and f) display slower diffusion rates.

#### Fewer SURF-TG

In the TG-rich LD simulations, SURF-TG levels decrease upon MLX binding, dropping to around 1.5% of the monolayer, while the opposite leaflet remains unchanged at 3 to 6%. This trend was consistent across all systems. SURF-TG abundance is known to correlate with applied surface tension (25, 37); under expansion, more TGs enter the monolayer to relieve tension, thereby promoting defect formation and enabling recruitment of proteins like CCTα (25). In contrast, binding of AH-containing proteins like MLX appears to displace PL headgroups, reduce local defects, and expel SURF-TGs into the core. This protein-induced shift in monolayer composition may affect the binding of other nearby proteins.

#### Decreased order below, increased order surrounding

PL tail-order parameters measure how ordered lipid tails are along their length, with C1 located near the glycerol backbone and C16/C18 at the tail end. Focusing on the POPC sn1 tail, association of MLX-DCD clearly increases the net order relative to the leaflet without protein (Figure 7e). This is also true for POPC sn2 and DOPE sn2 (Figure S16a, b). It is not the case, however, directly under the protein—PLs within 3 nm of MLX-DCD showed significantly decreased order as the tails fill the space the headgroups have been pushed away from (Figure S16d). Rather, lipids located more than 1 nm away from the protein explain the net higher order compared to protein-free membranes. This pattern was consistent in the DCD-double and DCD-big (Figure S16c, e) systems. Thus, the ordering effect does not appear to be an artifact of system size, as it persisted in the enlarged DCD-big system, nor of altered surface tension due to one leaflet accommodating the protein while the other does not, as the results are identical in DCD-double system. However, the effect does gradually weaken with distance, becoming minimal greater than 7 nm from the protein in the DCD-big system (Figure S16e). This pattern suggests that protein association locally disrupts lipid order while simultaneously enhancing order in the surrounding membrane. In real membrane systems the extent of increased packing order will likely depend on the membrane’s composition and density of associated proteins, with more crowded or rigid membranes being more affected than those capable of accommodating local stress through expansion.

#### Dynamics and Fluidity Analysis

To analyze the dynamic fluidity of the monolayer the mean square displacement (MSD) of PLs was evaluated in the DCD-big system. We found that the MSD for the lipids was larger in the leaflet without associated proteins (Figure 7f, S17). This was further corroborated by the lateral diffusion coefficients (Table S1), indicating that protein association reduces the fluidity of membrane PLs. These findings collectively demonstrate that the association of MLX-DCD with LDs not only changes the defect composition but also influences the ordering and dynamics of the surrounding membrane.

These findings align with broader experimental evidence that demonstrated AH binding to lipid bilayers can enhance adjacent leaflet order, potentially influencing membrane curvature and scission processes (68, 69). Our study extends such protein-induced surface effects to LD monolayers. Such altered properties could influence the co-localization of other MLX proteins, which could play an important role in regulating dimerization and dissociation. Alternatively, they could influence the proximal association of other proteins requiring either large packing defects or a direct interaction with surface-active TG.

## Conclusions

This work demonstrates that MLX specifically targets TG-rich lipid droplets by recognizing and embedding into shallow PL-acyl chain packing defects, distinguishing its mechanism from other CYToLD proteins that rely on deeper insertion into more pronounced TG-induced membrane defects (25, 34). Focusing on two sequences provides insight into how the shorter hairpin retains the ability to bind defect-rich membranes like the ER, while the longer DCD requires larger defects only accessible on the LD surface. The hairpin also serves as a model system offering insight into the full association mechanism of a CyTOLD-like AH containing protein. We find that neighboring residues first catch PL headgroups (Arg, Lys, Glu) and then dive into initial defects (Leu, Tyr, Phe). Multiple catch-dive pairs at opposite ends of helices can re-catch PLs if association is lost and anchor the helix once both sides form interactions. Following this, hydrophobic residues embed into the membrane as sufficient defects form. In addition, interhelical interactions that form upon binding disfavor dissociation, enhancing both affinity and retention on the LD surface. Collectively, these findings highlight how kinetic selection could influence protein selectivity in the LD proteome, wherein proteins with longer-lived catch-dive phases are more likely to stay associated while sufficient defects form for full embedding.

Turning to the DCD we find similar interactions wherein larger defects are required for full association, thereby explaining its strict selectivity for TG-rich LDs. Although the DCD structure folds in on itself in solution, it opens to a bound conformation that closely resembles that found in the MLX-DCD dimer interface. This structural mimicry suggests a shared interface that enables MLX to toggle between transcriptional regulation and LD localization depending on the cellular context. Exclusion from SE-rich LDs again stems from the relative absence of large packing defects, a feature likely common to other CYToLD proteins that fail to associate with SE-LDs. This selectivity links LD composition to MLX localization, coupling the lipid metabolic state with the regulation of glucose-responsive gene expression. Future studies could explore how tethering MLX or MYC-like proteins to the MLX-hairpin or DCD may shift regulatory outcomes.

Lastly, the altered surface properties near the bound MLX hairpin and MLX DCD, including reduced packing defect size, increased lipid tail order, and decreased lipid mobility, suggest that membrane remodeling occurs upon protein binding. These changes may contribute to form of protein-surface-protein allostery, influencing which proteins can subsequently associate with LDs and shaping patterns of colocalization on their surfaces.

## Acknowledgements

The authors are grateful for helpful discussions with Robert V Farese and Tobias C. Walther. The work was supported by the National Science Foundation under Grant No. (2341008) and the computational resources provided by Expanse through the San Diego Supercomputer through the Center Advanced Cyberinfrastructure Coordination Ecosystem (ACCESS) program (allocation MCB200018) supported by NSF (grant nos. 2138259, 2138286, 2138307, 2137603, and 2138296), as well as the Center for High-Performance Computing (CHPC) at the University of Utah. These resources greatly aided in the completion of this research.

## Author contributions

R.J.B and J.M.J.S designed the research. R.J.B. ran the simulations and analyzed the results. J.M.J.S directed the research. Both authors wrote the manuscript and agreed on the final version.

