## Supplementary Information for "Mechanistic Insights into the Selective Targeting of MLX to Triacylglycerol-Rich Lipid Droplets"

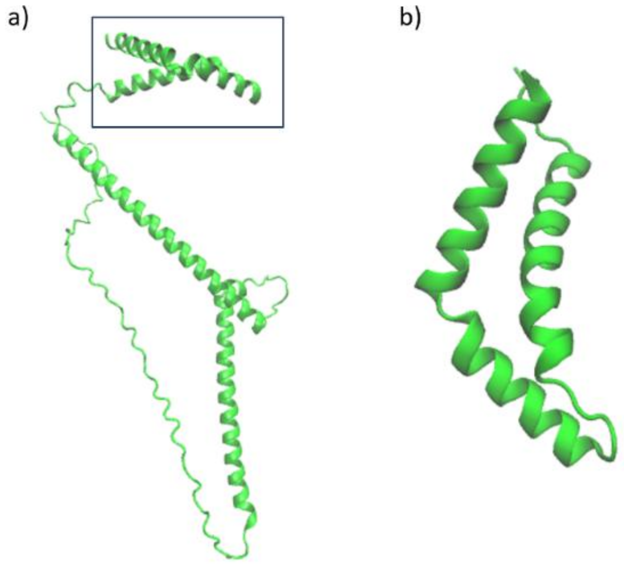


SI Figure 1. a) Predicted MLX monomer structure (DCD boxed) and b) corresponding MLX-DCD only viewed from above.

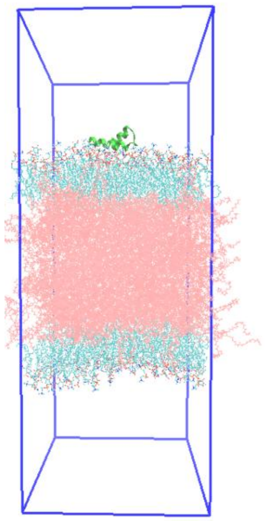

SI Figure 2. A LD trilayer simulation box setup with the PLs colored in cyan, TG in pink, and protein in green. Water is omitted for clarity.

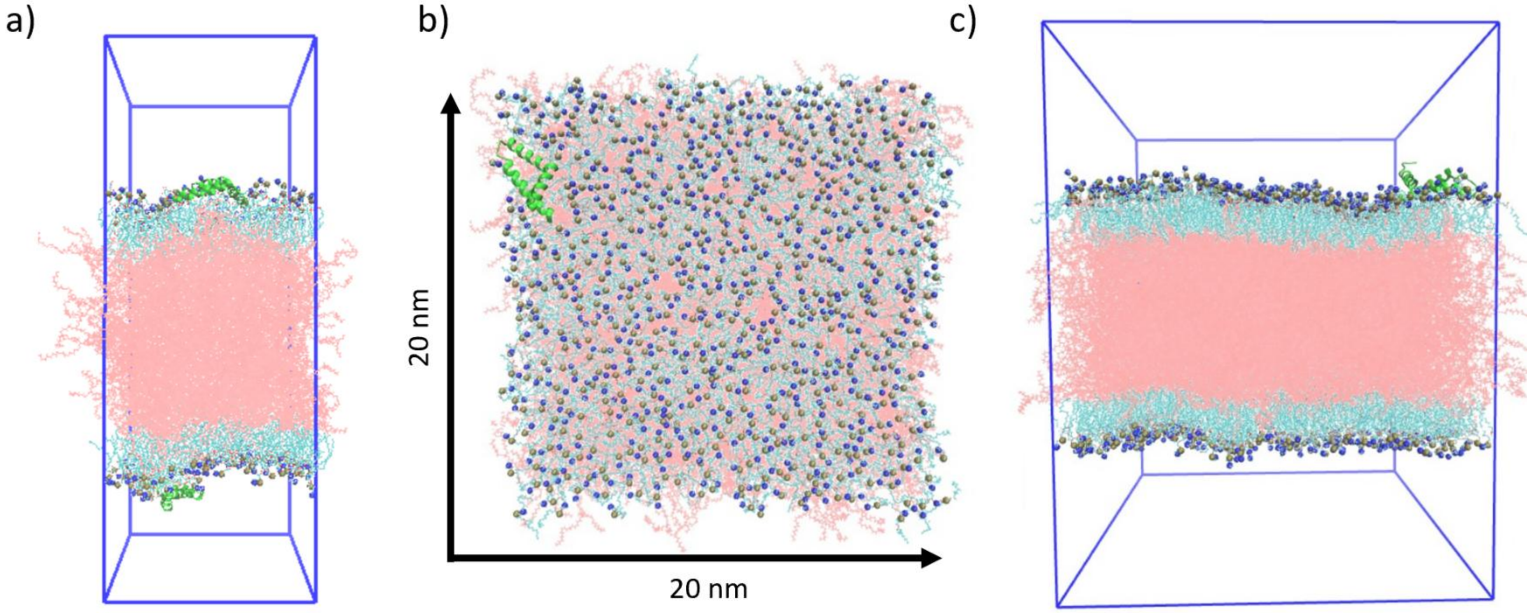


SI Figure 3. a) The DCD-double system with a protein on each leaflet. b) DCD-big has 4 times the surface area than the regular DCD LD system. c) Side view of DCD-big.

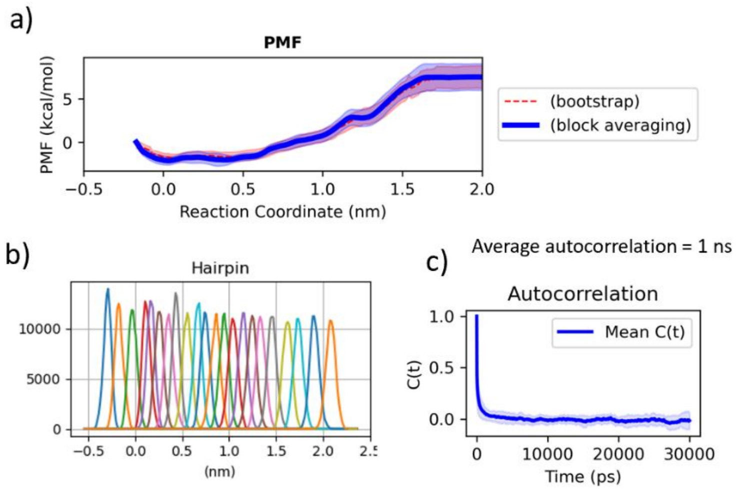


SI Figure 4. Example analysis taken from the initial 50:50 TG:SE MLX-hairpin association. a) Block averaging versus bootstrapping demonstrates similar error. b) Example histograms from hairpin binding to the LD demonstrate adequate overlap. c) Average autocorrelation for the CV per window averages 1 ns.

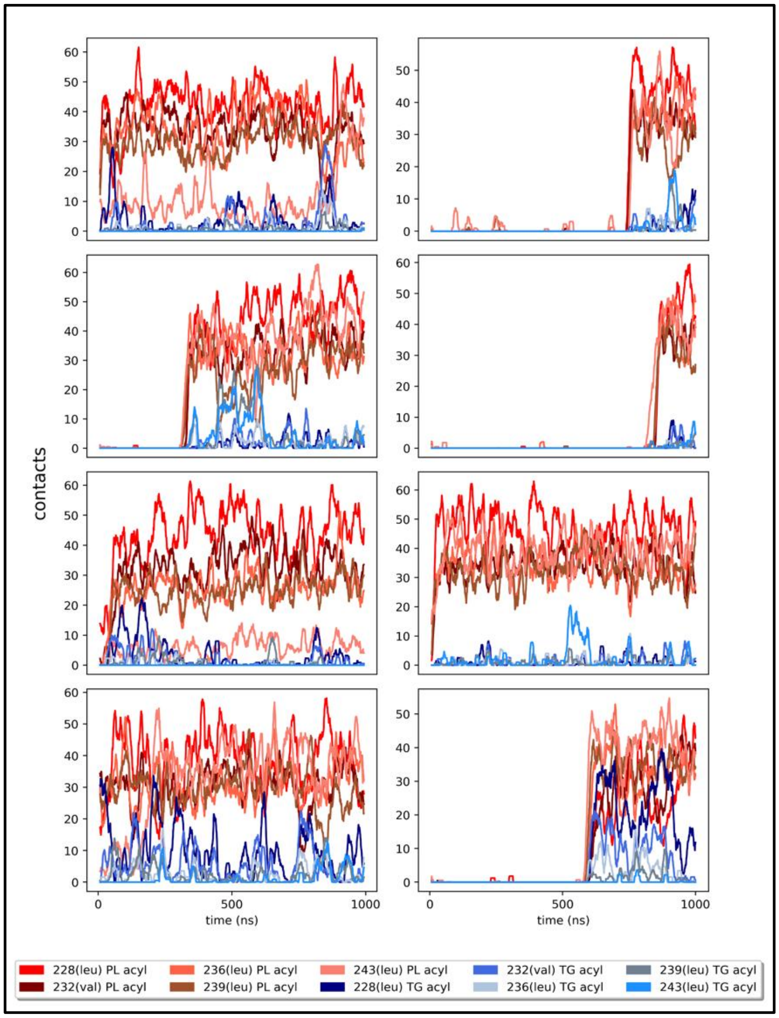


SI Figure 5. The 8 binding replicas of MLX with TG-LDs demonstrate that there are numerous contacts with the PL-acyl tails, as opposed to the TG molecules.

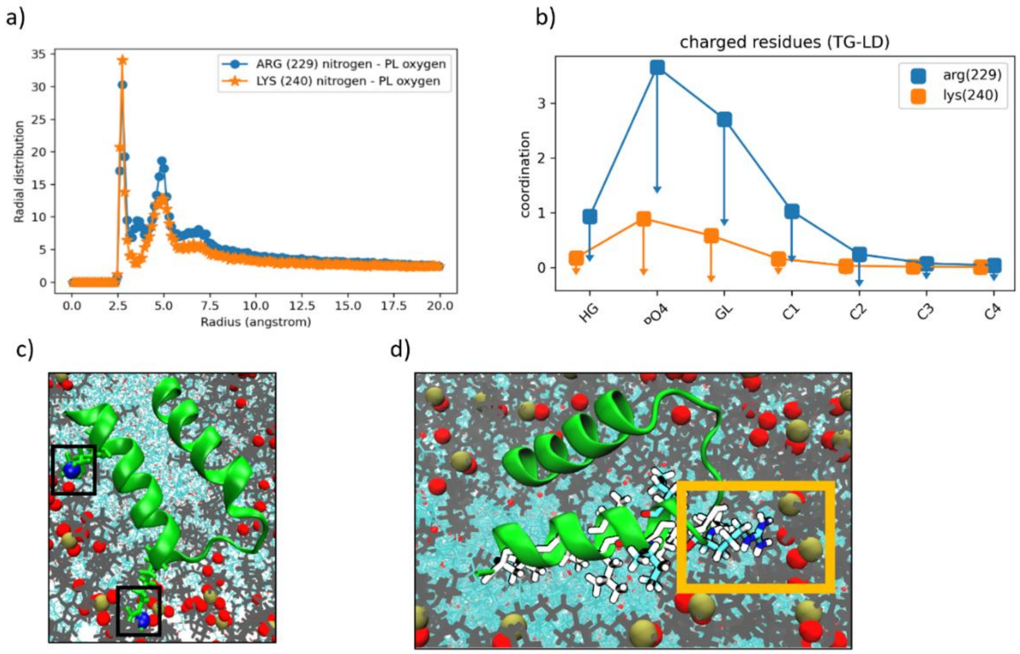


SI Figure 6. a) Radial distribution function (RDF) of Arg229 and Lys240 sidechains interacting with the PL-oxygens of headgroup. b) Coordination of Arg229 and Lys250 interacting with PLs highlighting the PL PO4 group as the dominant target. c) Arg229 and Lys240 (in black boxes) are on the polar side of the AH, interacting with PL-headgroups. d) Arg229 plays a critical role in not only 'catching', but also 'anchors' the AH using PL-headgroup interactions.

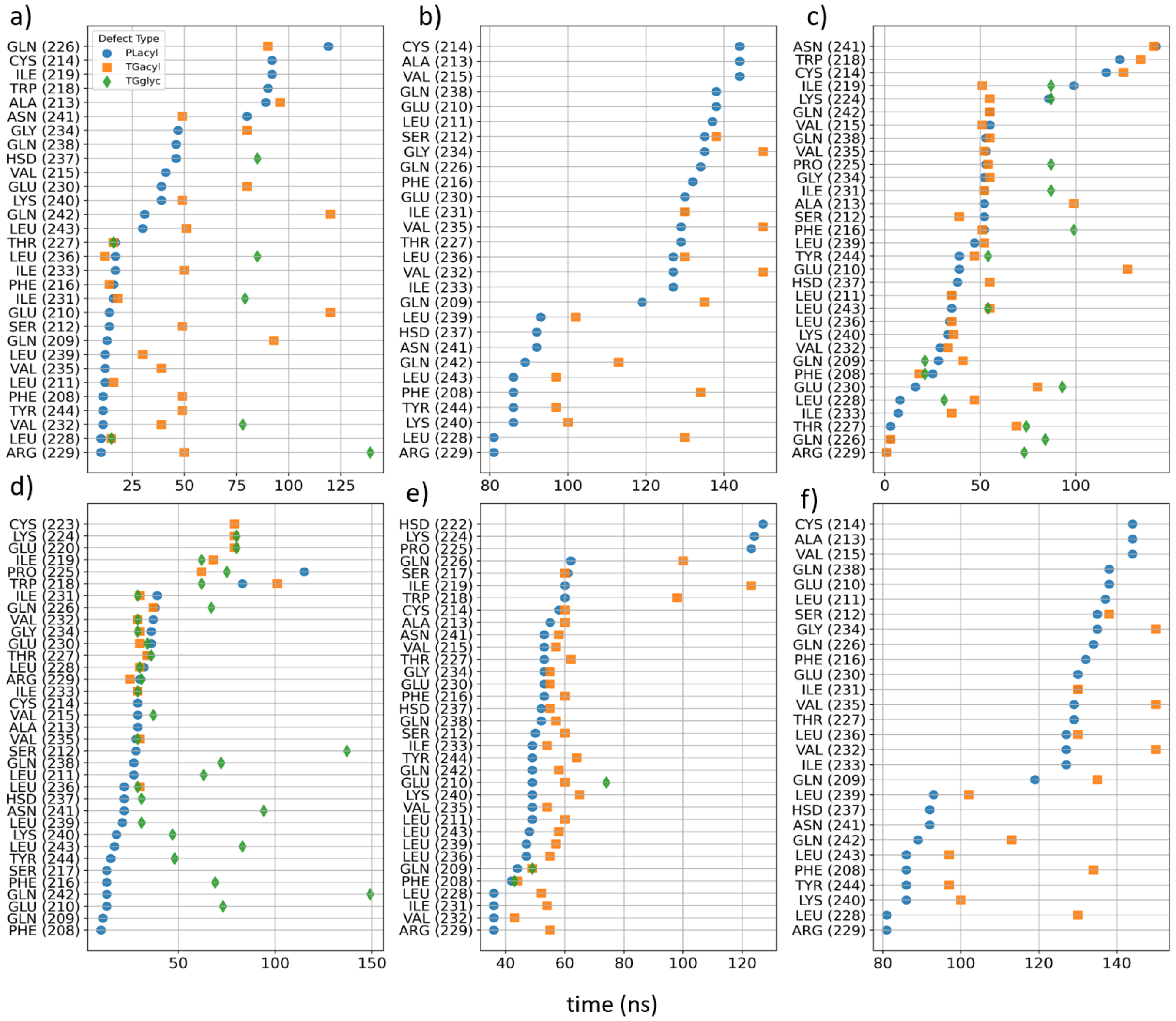


SI Figure 7. a-f) Time resolved defect interactions allow for tracking the specific timeline of how MLX binds to LDs. X-axis represents time, while the blue, orange, and green dots represent residue interactions with PL-acyl, TG-acyl, and TG-glycerol defects, respectively.

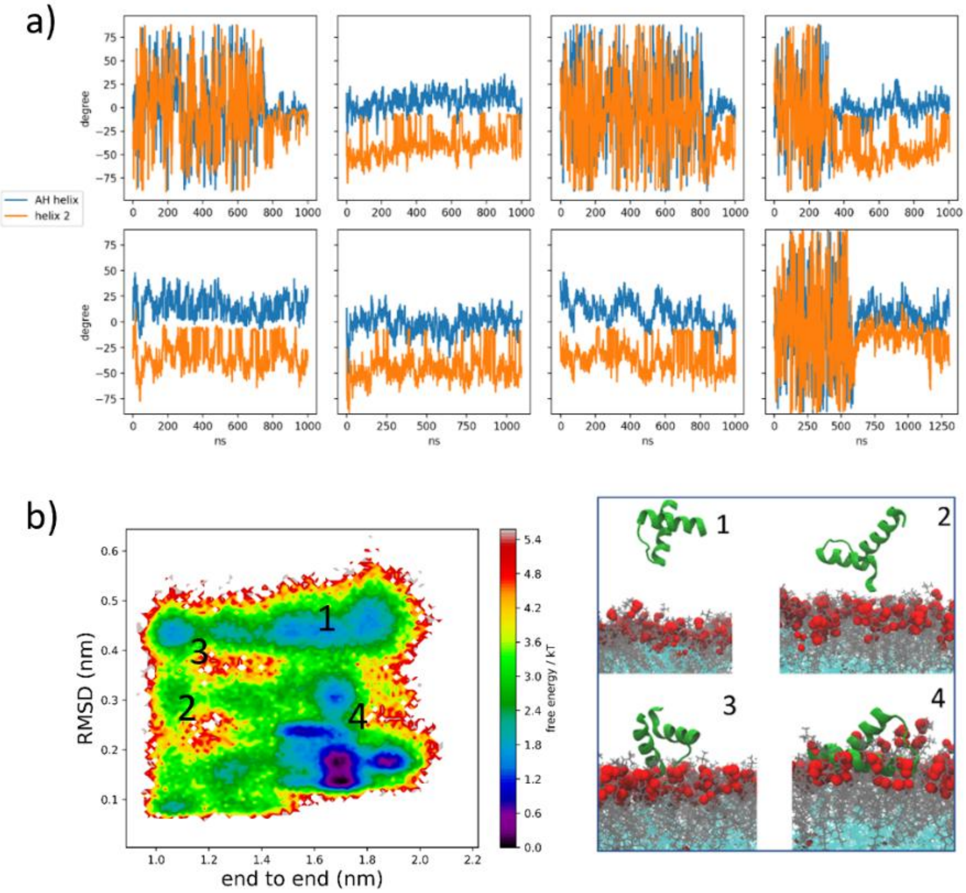


SI Figure 8. a) Tilt angles of the 8 MLX-hairpin replicas that bound to the TG-LD. The AH always tilts slightly upwards, and the helix-2 always tilts down. b) Free energy projection of the RMSD vs end-to-end protein distance.

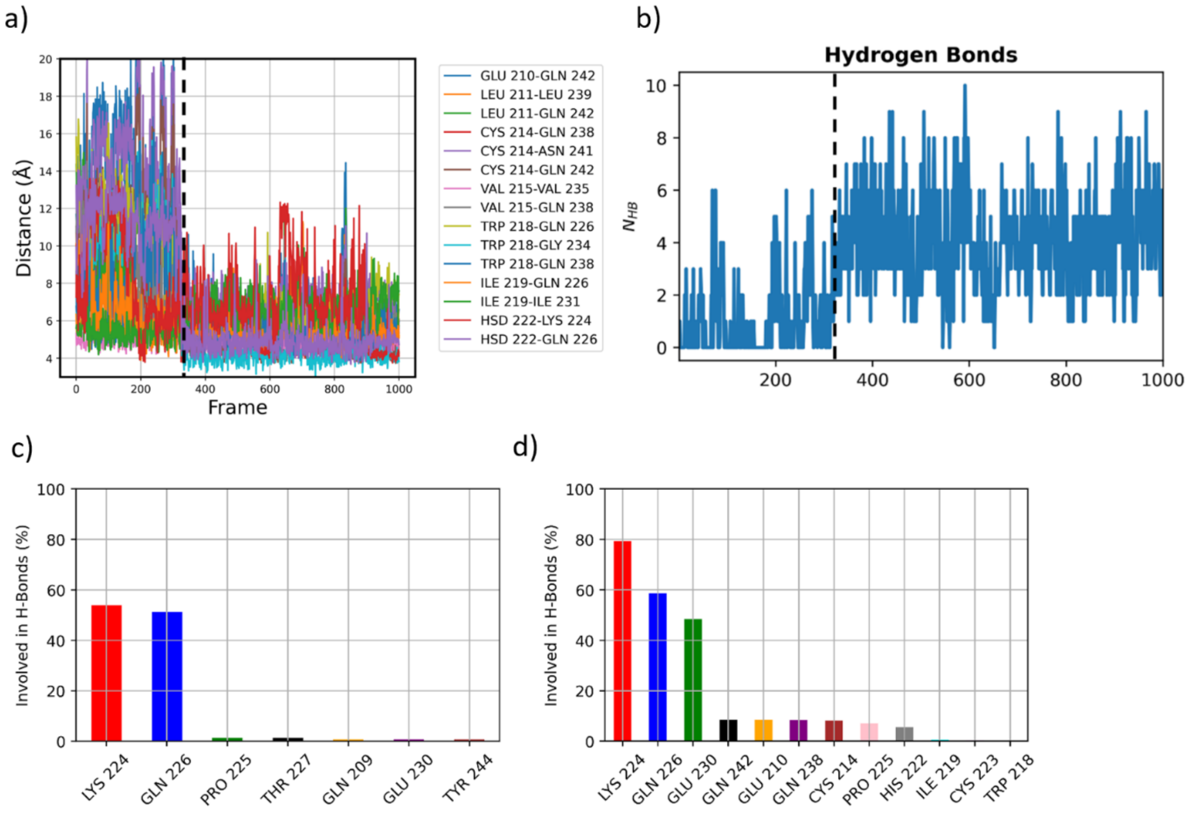


SI Figure 9. a) Distances between residues of the two helices of MLX when unbound and bound. The dotted black line refers to the moment of association to the LD monolayer. b) The number of hydrogen-bonds doubles when in the bound conformation. c) Hydrogen-bond types before association, and d) after association.

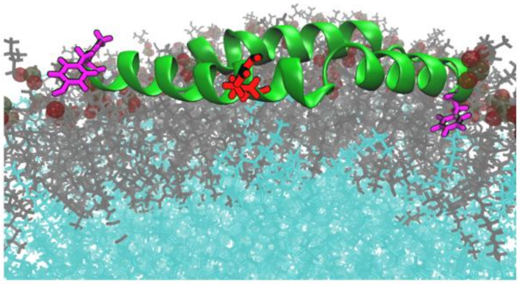

SI Figure 10. The catch and dive residues Arg228/Leu229 (center), Lys240/Tyr244 (left), and Glu210/Phe208 (right) are shown in red and magenta, respectively, interacting with the LD monolayer in a similar fashion as the MLX-hairpin.

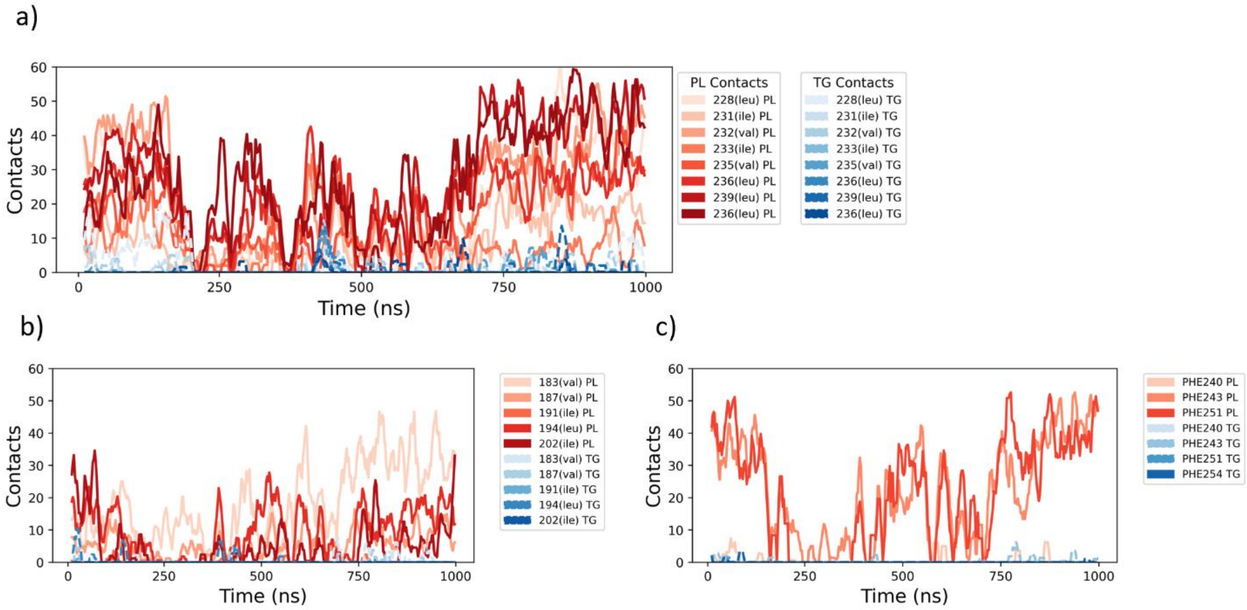


SI Figure 11. a) The hydrophobic residues of AH2 make numerous interactions with PL-acyl defects. b) The hydrophobic residues of AH2 also create numerous interactions, along with c) the Phe residues of the DCD.

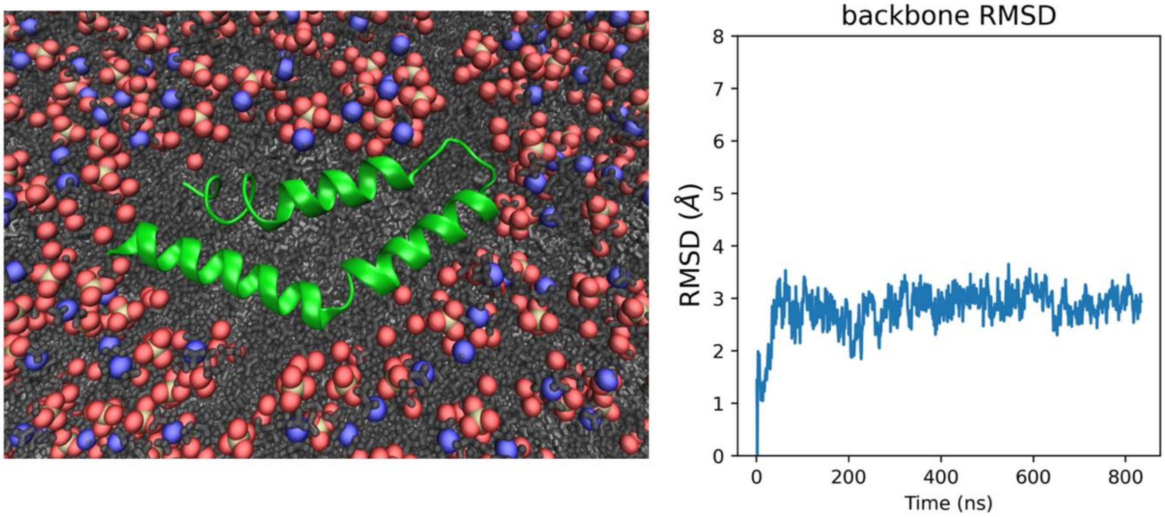


SI Figure 12. Top-down view of MLX-DCD bound the TG-LD (left), which has stable backbone RMSD upon LD association (right).

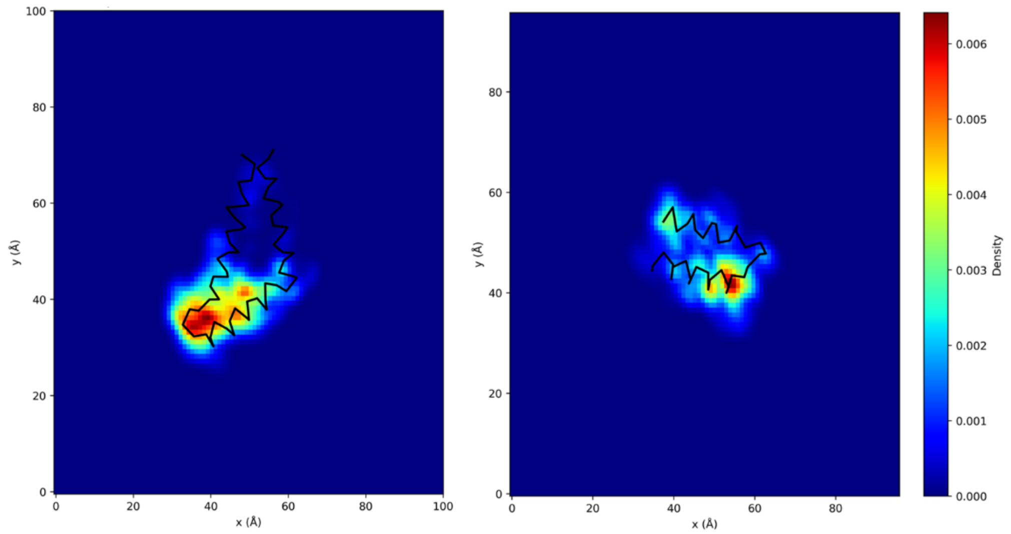


SI Figure 13. Defect size of MLX DCD (left), and MLX-hairpin (right).

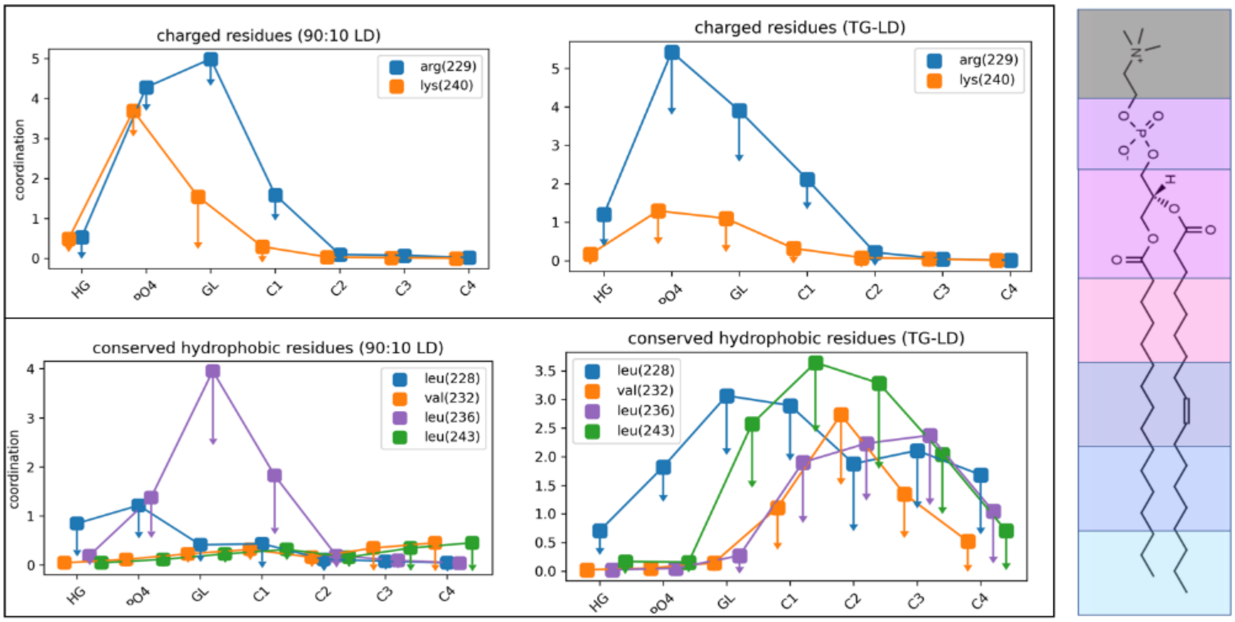


SI Figure 14. Coordination analysis extracted from the umbrella sampling frames in which the hairpin was bound to the LD. While the charged residues in both the 90:10 and pure-TG systems are interacting with the headgroups, the hydrophobic residues are able to penetrate deep into PL-acyl defects in the pure-TG LD.

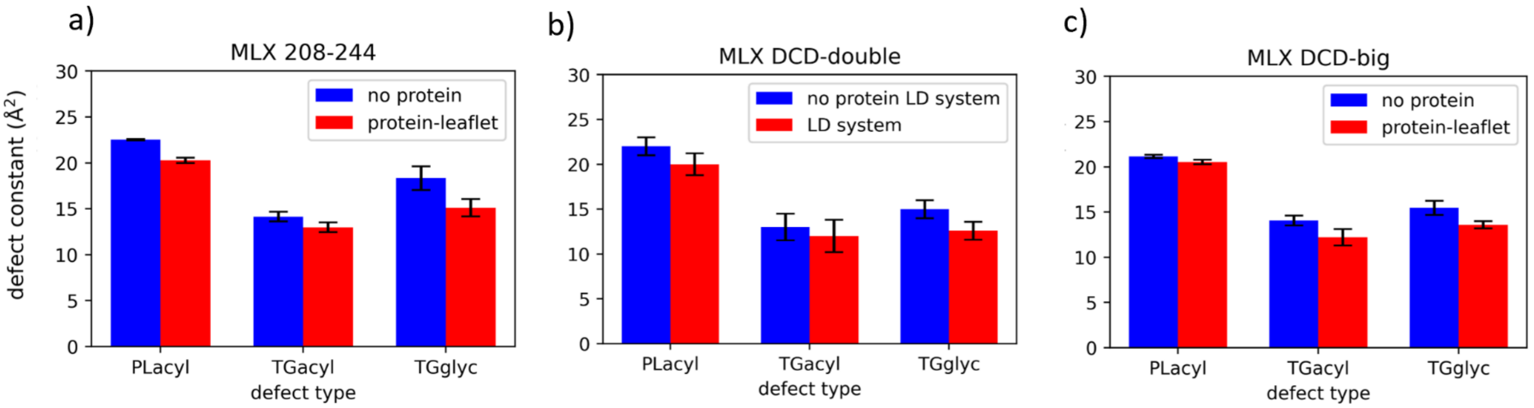


SI Figure 15. Packing defect constants for a) MLX-hairpin, b) DCD-double, and c) DCD-big. For the DCD-double system, a reference LD with no protein was used to compare against the protein-embedded LD system.

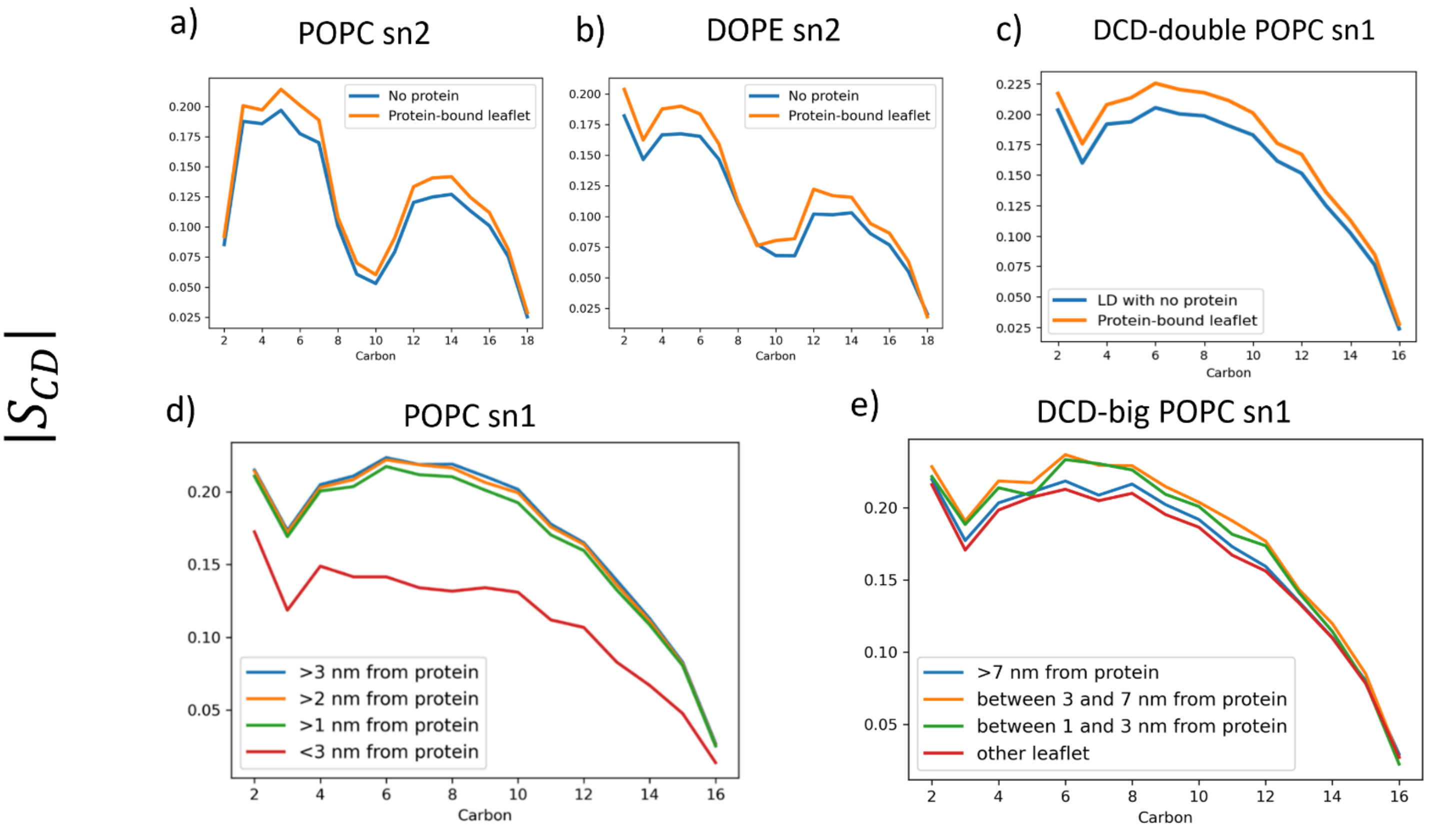


SI Figure 16. Tail order parameters for (a) POPC sn2 and (b) DOPE sn2 in the MLX-DCD TG-rich LD. (c) The tail order for POPC sn1 in the DCD-double system is nearly identical to the MLX-DCD (Figure 7e), with blue representing a reference system with no protein on either leaflet. d) Tail order underneath the MLX-DCD protein (< 3nm) is substantially decreased, while tails >1 nm away are consistently more ordered. e) As you move further away (> 7nm) in the MLX DCD-big system the increased ordering dissipates.

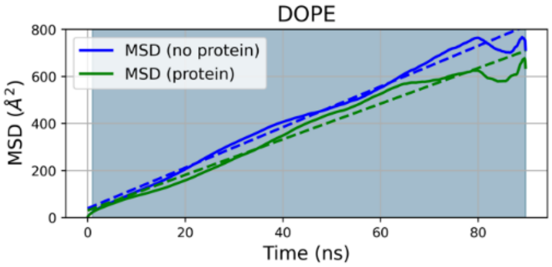


SI Figure 17. MSD for DOPE PLs.

| | |
|---|---|
| **POPC no protein** | $21.6 \times 10^{-8}\ \frac{cm^2}{s}$ |
| **POPC protein** | $16.1 \times 10^{-8}\ \frac{cm^2}{s}$ |
| **DOPE no protein** | $21.7 \times 10^{-8}\ \frac{cm^2}{s}$ |
| **DOPE protein** | $18.9 \times 10^{-8}\ \frac{cm^2}{s}$ |
| **SAPI no protein** | $28.8 \times 10^{-8}\ \frac{cm^2}{s}$ |
| **SAPI protein** | $18.9 \times 10^{-8}\ \frac{cm^2}{s}$ |

SI table 1. MSD values for the various PLs of the DCD-LD system

### Supplementary Methods

#### Structure determination

In the absence of experimental structural information, a variety of tools were used to predict the structures of MLX-hairpin and MLX-DCD. MLX-hairpin was initially folded using Rosetta Ab Initio[1,2], with a substantial replica count of 50,000. Initial broad conformational space exploration was followed by clustering to identify promising structures, which were then refined through targeted design protocols. This strategy ensured comprehensive coverage and accuracy in predicting protein folds and interactions within computational constraints. From this set, we used the Rosetta scoring function to determine 10 most stable structures based off the free energy exhibited notable structural similarity.[3,4] As protein folding methods evolved, we turned to the highly reliable AlphaFold2 and RoseTTAFold.[5–7] The following parameters were used for AlphaFold2: *num_relax=1, template_mode=none, msa_mode=mmseqs2, template_mode=none, pair_mode=unpaired_paired, modetl_type = auto, num_recycles=24, recyle_early_stop_tolerance=auto, relax_max_iterations=200,* and for RoseTTAFold: *msa_method=mmseqs2, paire_mode=unpaired_paired, num_recycles=24, max_msa=256*

Additionally, the structure predictions for the whole structure of MLX is available at the AlphaFold protein structure database: https://alphafold.ebi.ac.uk/entry/Q9UH92, Where the DCD and MLX-hairpin domain receive a very high *per-residue model confidence score* pLDDT score.[8]

The predictions of all models once again yielded similar structures, with an AH-kink-AH structure. We proceeded with the initial structure folded through Rosetta ab initio as it had already been equilibrated. MLX-DCD was predicted with both AlphaFold2 and RoseTTAFold, and again all the structures agreed in alignment, with a structure composed of 3 helices that were folded in a planar triangular structure, to shield hydrophobic residues from water (SI Figure 1).
